# Autophagy promotes photomorphogenesis during seedling development in Arabidopsis in carbon limiting conditions

**DOI:** 10.1101/2021.03.25.437007

**Authors:** Akila Wijerathna-Yapa, Santiago Signorelli, Ricarda Fenske, Diep R. Ganguly, Elke Stroeher, Lei Li, Barry J. Pogson, Owen Duncan, A. Harvey Millar

## Abstract

Autophagy is a conserved catabolic process that plays an essential role under nutrient starvation condition and influences different developmental processes. We observed that seedlings of autophagy mutants (*atg2*, *atg5*, *atg7,* and *atg9*) germinated in the dark showed delayed chloroplast development following illumination. The delayed chloroplast development was characterized by a decrease in photosynthetic and chlorophyll biosynthetic proteins, lower chlorophyll content, reduced chloroplast size, and increased levels of proteins involved in lipid biosynthesis. Confirming the biological impact of these differences, photosynthetic performance was impaired in autophagy mutants 12h post illumination. We investigated if the delayed chloroplast development could be explained by lower lipid import to the chloroplast or lower triglyceride (TAG) turnover. We observed that the limitations in the chloroplast lipid import imposed by *trigalactosyldiacylglycerol1* are unlikely to explain the delay in photomorphogenesis. However, we found that lower TAG mobility in the triacylglycerol lipase mutant *sugardependent1* significantly affected photomorphogenesis. Moreover, we showed that lower levels of carbon resources exacerbated the delay in photomorphogenesis whereas higher levels of carbon resources had an opposite effect. This work provides evidence that autophagic process operate during de-etiolation in a manner that contributes to photomorphogenesis through increasing lipid turnover to physically or energetically sustain photomorphogenesis.

## Introduction

Autophagy is a catabolic cellular process in which cytoplasmic content and organelles are degraded in lytic vacuoles (micro- and macro-autophagy) or in the cytosol (mega-autophagy). In the latter, the degradation of cellular content is caused by the release of vacuolar lytic content in the cytosol leading to cell death. However, micro- and macro- autophagy are processes contributing to cell maintenance and have been proved to be relevant in plant cell biology (Bassham 2007; Masclaux-Daubresse et al. 2017). Autophagy is performed by a series of autophagy-related (ATG) proteins which constitutes the autophagy core machinery, involved in the formation of autophagosomes and their fusion to lytic vacuoles (Marshall and Vierstra 2018).

Autophagic activity is known to be induced under certain conditions such as carbon and nitrogen starvation and to promote recycling of these elements (Avin-Wittenberg et al. 2015; Di Berardino et al. 2018; Luo et al. 2019). Due to this capacity, autophagy has been reported to be a key factor for developmental processes such as seed filling and maturation (Chung et al. 2009; Guiboileau et al. 2012; Janse van Rensburg et al. 2019). Autophagy can also be induced under a wide range of abiotic and biotic stress conditions, and its activity is known to contribute to plant stress tolerance (Avin-Wittenberg 2018; Signorelli et al. 2019).

In the past few years, sub-types of autophagy that influence specific cellular organelles have been reported, such as the mitophagy, chlorophagy, and pexophagy, among others, for the specific removal of mitochondria, chloroplasts and peroxisomes, respectively (Li et al. 2014; Young and Bartel 2016; Izumi et al. 2017). During seed germination and the early stages of seedling development, biogenesis of different organelles occurs and each play a role in metabolic transitions involved in utilising carbon resources from the seed and then from the atmosphere to sustain plant growth (Law et al. 2014; Pogson et al. 2015). During leaf senescence, selective degradation of organelles is reported to occur, linking mitophagy and chlorophagy (Izumi et al. 2010; Ishida et al. 2014; Broda et al. 2018).

In this work, first we used multiple reaction monitoring (MRM)-based targeted proteomics to assess the abundance of selected organelle proteins during early seedling development (i.e., seed reserve mobilisation), during dark-to-light transition of etiolated seedlings (i.e., photomorphogenesis) and light to dark transition of mature leaves (i.e., dark induced senescence) in different autophagy mutants and wild-type (WT). Our results highlight a chloroplast biogenesis delay in dark to light transition of etiolated seedling in autophagy mutants revealing new links between autophagy and i) the accumulation of photosynthetic machinery during photomorphogenesis, ii) the carbon status of seedlings and iii) plastid-dependent lipid mobilisation.

## Results

### Organelle abundance is selectively affected in autophagy mutants under different developmental transitions

We defined three different developmental transitions, early seedling development (T1), dark-to-light greening of seedlings (T2), and light-to-dark senescence of leaves (T3), to investigate how organelle development processes are affected in autophagy mutants. To follow the differences in organelle development, we measured the abundance of organelle specific marker proteins in each developmental transition using a targeted proteomics (MRM) strategy, established in a previous study (Hooper et al. 2017). The protein markers used included key enzymes from the chloroplast, mitochondria, peroxisome, ER, Golgi, and plasma membrane, as well as subunits of the cytosolic ribosome. Among the studied markers, all the chloroplast proteins (PHOTOSYSTEM II SUBUNIT P-1, CHLOROPLAST RNA BINDING, LIGHT HARVESTING COMPLEX B6 OF PHOTOSYSTEM II and GERANYLGERANYL DIPHOSPHATE REDUCTASE) increased in abundance one day after the dark to light transition (**Figure 1A-B**), most were also induced in the early seedling development transition, but none of them changed in the light to dark senescence transition (**Figure S1**). When we analysed these organelle protein markers across the three developmental transitions in *atg2*, *atg5*, *atg7* and *atg9*, we observed all four *atg* lines showed a delay in the increase of chloroplast markers during dark to light transition. Chloroplast markers in most of the autophagy mutants were less abundant than WT at one day after light treatment (**Figure 1C**). Unlike the chloroplast markers, other organelles’ protein markers showed no or inconsistent change during the dark to light transition (**Figure S2**). In the other two developmental transitions, no consistent differences were observed between *atg* lines and WT for any set of organelle markers (**Supp. File 1 and 2**).

**Figure 1.**
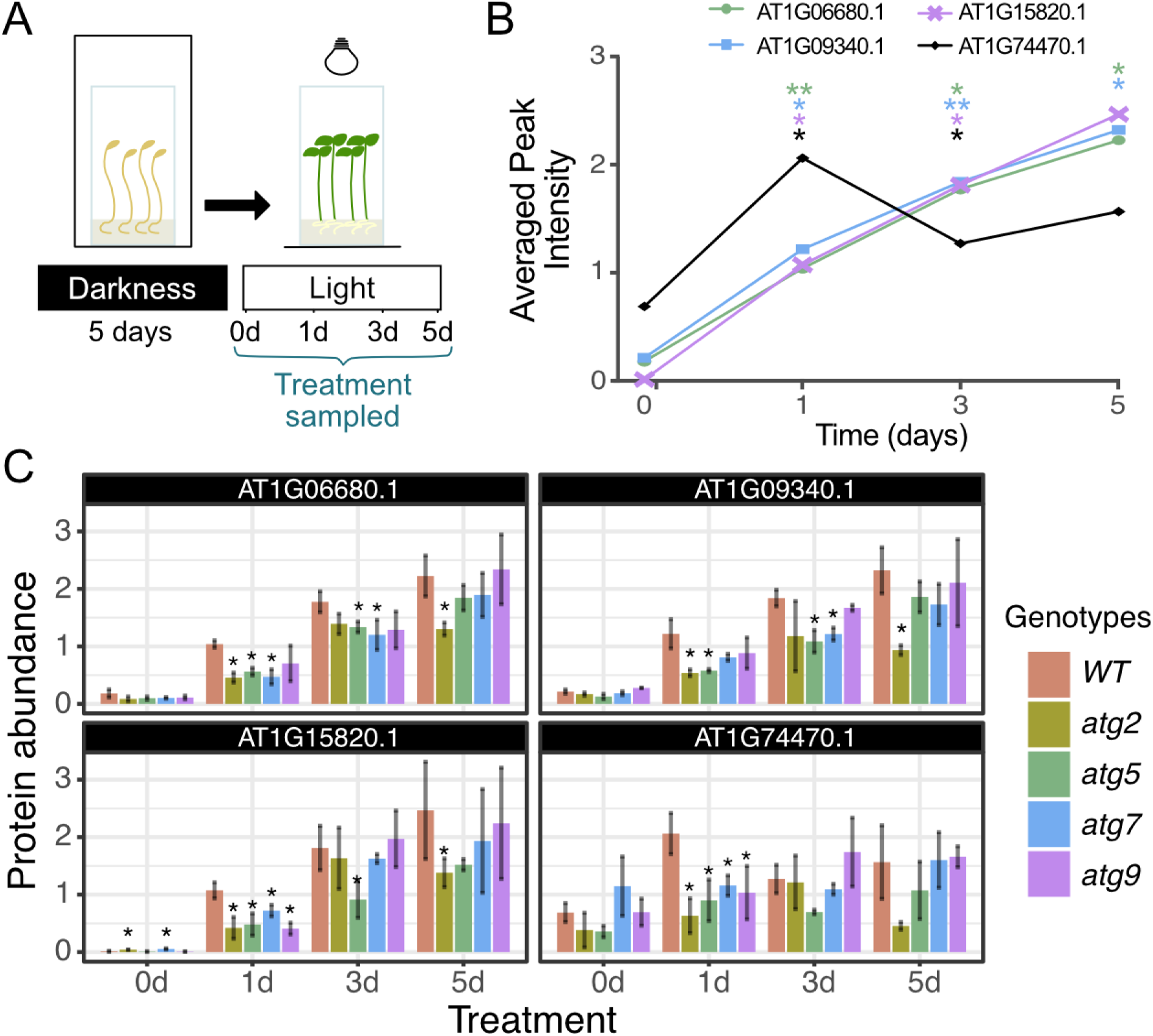
The dark to light transition for seedling greening is affected in atg lines and characterized by less abundant chloroplast markers. **A.** Experimental design showing the dark to light transition. **B.** Abundance of plastid MRM biomarkers, viz. AT1G06680.1 (PHOTOSYSTEM II SUBUNIT P-1), AT1G09340.1 (CHLOROPLAST RNA BINDING), AT1G15820.1 (LIGHT HARVESTING COMPLEX B6 OF PHOTOSYSTEM II), and AT1G74470.1 (GERANYLGERANYL DIPHOSPHATE REDUCTASE), during the dark to light transition in WT. The y- axis of the line graph represents averaged peak intensity of three biological samples (n=3) of these markers over the time course. Statistical analyses were performed by one-way ANOVA, followed by Tukey’s HSD test compared to the 0d time (*P < 0.05,). **C.** Abundance of plastid MRM biomarkers in WT Col-0 and autophagy mutants *atg2*, *atg5*, *atg7* and *atg9* during the dark to light transition (0, 1, 3 and 5 days). Means ± SE for n = 4 are shown. Asterisks denote statistically significant differences at the P < 0.05 level among treatment groups within a time-point.

### Altered greening and chloroplast development occur in autophagy mutants transferred from dark to light conditions

We selected the dark to light transition to have a further understanding about the role of autophagy during the photomorphogenesis. Because the most significant differences between WT and autophagy mutants were observed at early stages of light exposure (**Figure 1C**), for further experiments we focused on 0h, 12h and 24h after exposure to light (**Figure 2A**). At 0h all the seedlings were equally pale and presented a similar phenotype (**Figure 2B-C**); however, at 12h, WT plants were significantly greener than all the autophagy mutants based on visual inspection and calculation of a Green Index (GI) as outlined in the methods section (**Figure 2B-C**). At 24h the GI of seedlings did not increase further in WT, suggesting they already reached a maximum by 12h, whereas the autophagy mutants still increased their GI over that time period (**Figure 2B-C**). Light treatment also increased the total chlorophyll content measured in all the lines; however, WT plants had more chlorophyll than most autophagy mutants, especially at 12h (**Figure 2D**). Furthermore, we analysed the chlorophyll auto-fluorescence by confocal laser scanning microscopy as an indicator for presence and size of chloroplasts (**Figure 2E**). We observed that at 12h both the number of chloroplasts per area and chloroplast size were reduced in autophagy mutants, whereas by 24h most of the differences between WT and autophagy mutants were lost (**Figure 2F-G**).

**Figure 2.**
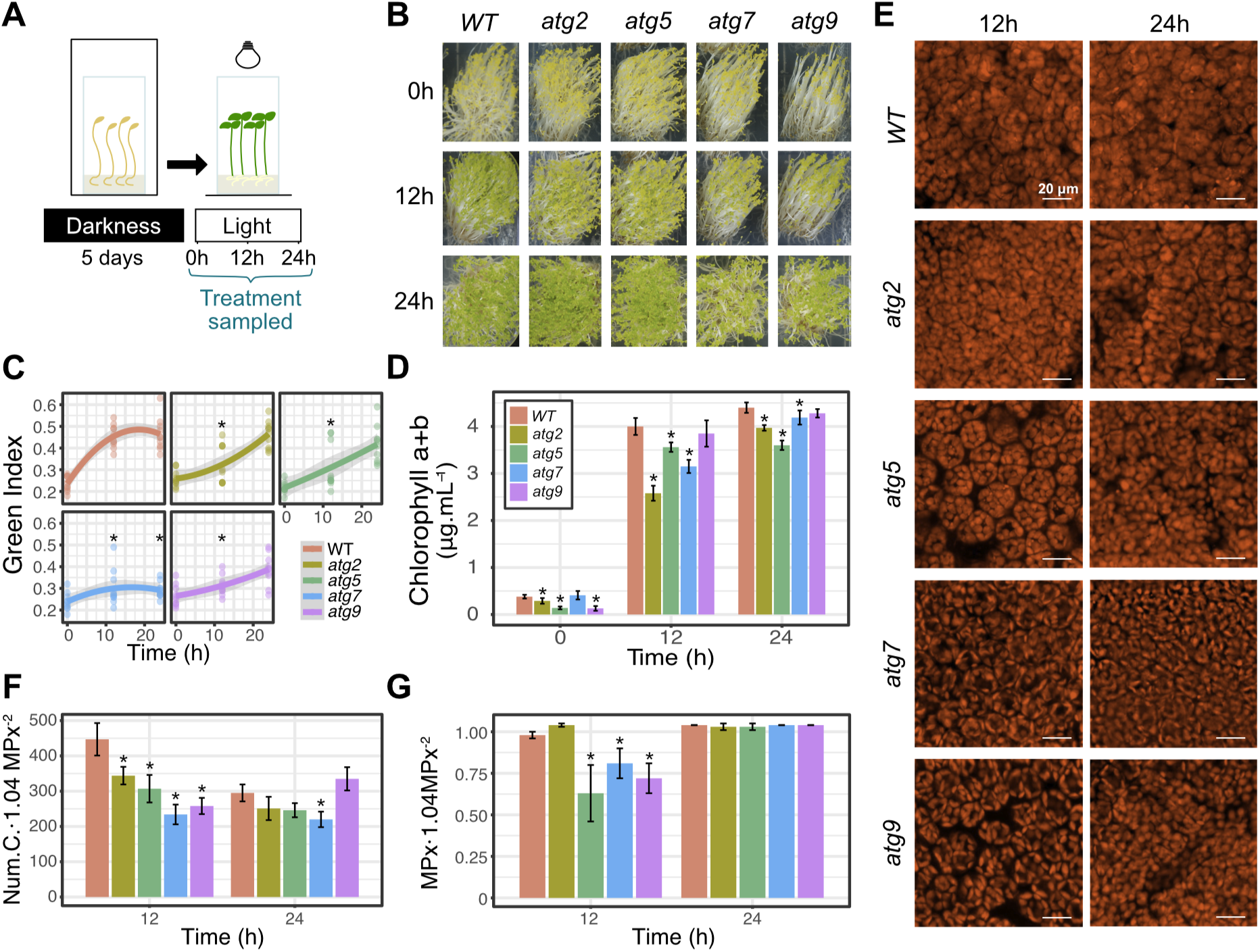
The dark to light transition in autophagy mutants is characterized by slower greening, lower chlorophyll content, and reduced number and size of chloroplast than the WT. **A.** Experimental design to study the dark to light transition for seedling greening. Seeds were germinated and grown on MS solid medium supplemented with 0.5% sucrose for 5 days in dark and then exposed to light for 0h, 12h and 24h. **B.** Phenotype of WT and autophagy mutants at 0h, 12h and 24h of light treatment. **C.** Green Index for WT and autophagy mutants at 0h, 12h and 24h of light treatment. The grey regression represents +/- 95% confidence level intervals for prediction for the 2-order polynomial regression. **D.** Chlorophyll content for WT and autophagy mutants at 0h, 12h and 24h of light treatment. (n= 4 independent biological replicates with three technical replicates per group). **E.** Chlorophyll autofluorescence of WT and autophagy mutants at 12h and 24h of light treatment. Autofluorescence was detected by CLSM at 640 nm using a excitation of 488 nm (scale bars: 20 μm). **F.** Number of plastids per area, and **G.** plastid area for WT and autophagy mutants at 12h and 24h of light treatment. Resolution size 0.07 μm/px, dimension of 1024x1024 px (equivalent to 1.04 MPx^2^) of autophagy mutants with control WT are shown (n= 5 independent biological replicates per group). Asterisks indicate P values ≤ 0.05 by two-way analysis of variance (ANOVA) and Tukey HSD multiple comparison test. Data are represented as mean ± S.D.

### Lower photosynthetic activity in de-etiolated seedlings of autophagy mutants

To understand if the differences observed between autophagy mutants and WT in terms of chlorophyll content and chloroplast development have physiological consequences we evaluated the photosynthetic activity by different methods. First, we measured the chlorophyll fluorescence related parameters, maximum quantum efficiency of photosystem II (F_V_/F_M_) and the quantum yield of non-regulated energy dissipation (**φ**NO). As observed in **Figure 3A** there is a clear contrast between F_V_/F_M_ at 12h and 24h for all lines further confirming that at 12h seedlings are not fully developed and have not reached their maximum photosynthetic capacity. At 12h, the F_V_/F_M_ was higher in WT than in the autophagy mutants (**Figure 3B**) indicating that WT plants were already more developed than autophagy mutants and thus made a more efficient use of incident light. However, at 24h F_V_/F_M_ of autophagy mutants was more similar to WT values, suggesting that autophagy mutants eventually attained normal development. Likewise, **φ**NO values in autophagy mutants were more different to WT values at 12h than at 24h, being always higher in autophagy mutants than WT values. We further investigated the relative apparent electron transport rate (ETR) parameter observing that in all lines the ETR values increased from 12h to 24h and in most cases the peak of ETR was shifted towards greater PAR from 12h to 24h (**Figure 3C**). This finding suggests a greater photosynthetic performance and chloroplast development was evident by 24h across all lines. However, WT ETRs were higher than ETR of autophagy mutants at both 12h and 24h and the WT peak was shifted to higher PAR values compared to autophagy mutants, which showed a more rapid decay at high light intensity (**Figure 3C**). Finally, we tested the oxygen evolution of seedlings as an indicator of photosynthetic activity and respiration, under light and dark conditions respectively. Respiration rates were similar over the period of 24h evaluated and among the different genotypes (**Figure S4 A**), however, photosynthetic activity increased over the same period and varied between genotypes (**Figure S4 B**) and so did the photosynthesis over respiration ratio (**Figure 3D**). The latter ratio is an indication of how photoautotrophic the seedlings were and thus of their developmental state. In particular, the photosynthesis over respiration ratio was similar between WT and autophagy mutants at 0h and 24h but significantly higher in WT than *atg5*, *atg7* and *atg9* at 12h (**Figure 3D**). Considering the different indicators for photosynthetic performance used here, we conclude that at 12h the autophagy mutants had lower photosynthetic performance than WT, but the differences were diminished by 24h.

**Figure 3.**
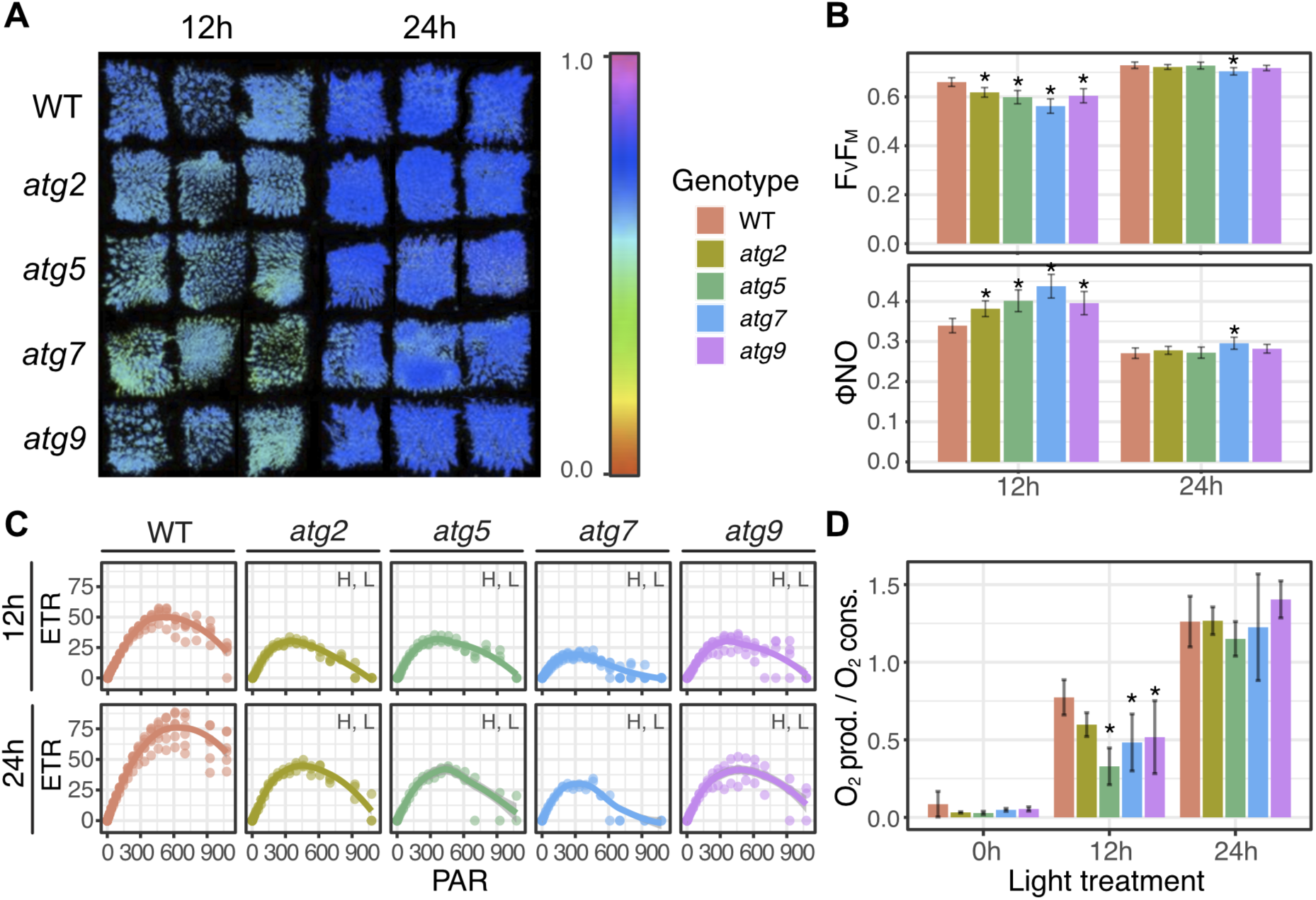
>Photosynthetic performance of autophagy mutants during the dark to light transition. **A.** F_V_/F_M_ images for WT and autophagy mutants at 12h and 24h. **B.** Analysis of F_V_/F_M_ and **φ**NO values for WT and autophagy mutants at 12h and 24h. **C.** Relative apparent electron transport rate (ETR) plots for WT and autophagy mutants at 12h and 24h. The grey regression represents +/- 95% confidence level intervals for prediction of the local regression. H and L, represent significantly higher values for WT with respect to autophagy mutants in terms of the height (H) and length (L) of the maximum ETR as explained in Figure S3. **D.** Ratio between photosynthetic activity and respiration for WT and autophagy mutants at 0h, 12h and 24h. Asterisks indicate statistically significant differences (P < 0.05) by Tukey’s HSD test compared to WT. Data are represented as mean ± S.D.

### Proteomic analysis of autophagy mutants’ photosynthesis machinery during de-etiolation

To better assess the differential chloroplast development in the autophagy mutants during the dark-light transition, we investigated the proteome of the autophagy mutants and WT plants at 0h, 12h and 24h by quantitative mass spectrometry, and selected plastid-localized proteins for further analysis (**Supp. File 3**). We identified differentially enriched plastid proteins (DEPP) and subsequently the enrichment of specific metabolic pathways and biological process. Compared to WT, at 0h, the autophagy mutants *atg2*, *atg5*, *atg7*, and *atg9* had lower abundance of 96, 83, 67, and 94 plastid proteins, respectively, and greater abundances of 32, 41, 47, and 25 plastid proteins, respectively. Among these DEPP, 51 lower abundance plastid proteins were common to all *atg* lines and 74 were common to at least three *atg* lines. Likewise, 11 more abundant plastid proteins were common to all *atg* lines and 29 were common to at least three *atg* lines (**Figure 4A**). Similar patterns were observed at 12h and 24h (**Figure 4A**). These data show a high consistency among the DEPP in the different autophagy mutants and across the timecourse (**Figure 4B**). In particular, we observed that 87 were more abundant with respect to WT in at least 3 *atg* lines in at least two time points. On the other hand, 34 were more abundant than in WT in at least 3 *atg* lines and at two time points. To explore these protein sets we analysed the GO biological process (GO BP) and KEGG pathway enrichment of this subset of consistent DEPP. The plastid proteins that were less abundant in at least 3 *atg* lines and at least two of the three times analysed showed an enrichment of proteins involved in photosynthesis and porphyrin and chlorophyll metabolism according to KEGG pathways, or an enrichment of chloroplast biogenesis, photosynthesis, and chlorophyll metabolism according to GO BP (**Figure 4C**). On the other hand, the more abundant plastid proteins showed an enrichment of fatty acid biosynthesis according to KEGG pathway or an enrichment of fatty acid biosynthesis and sugar catabolism according to GO BP (**Figure 4C**).

**Figure 4.**
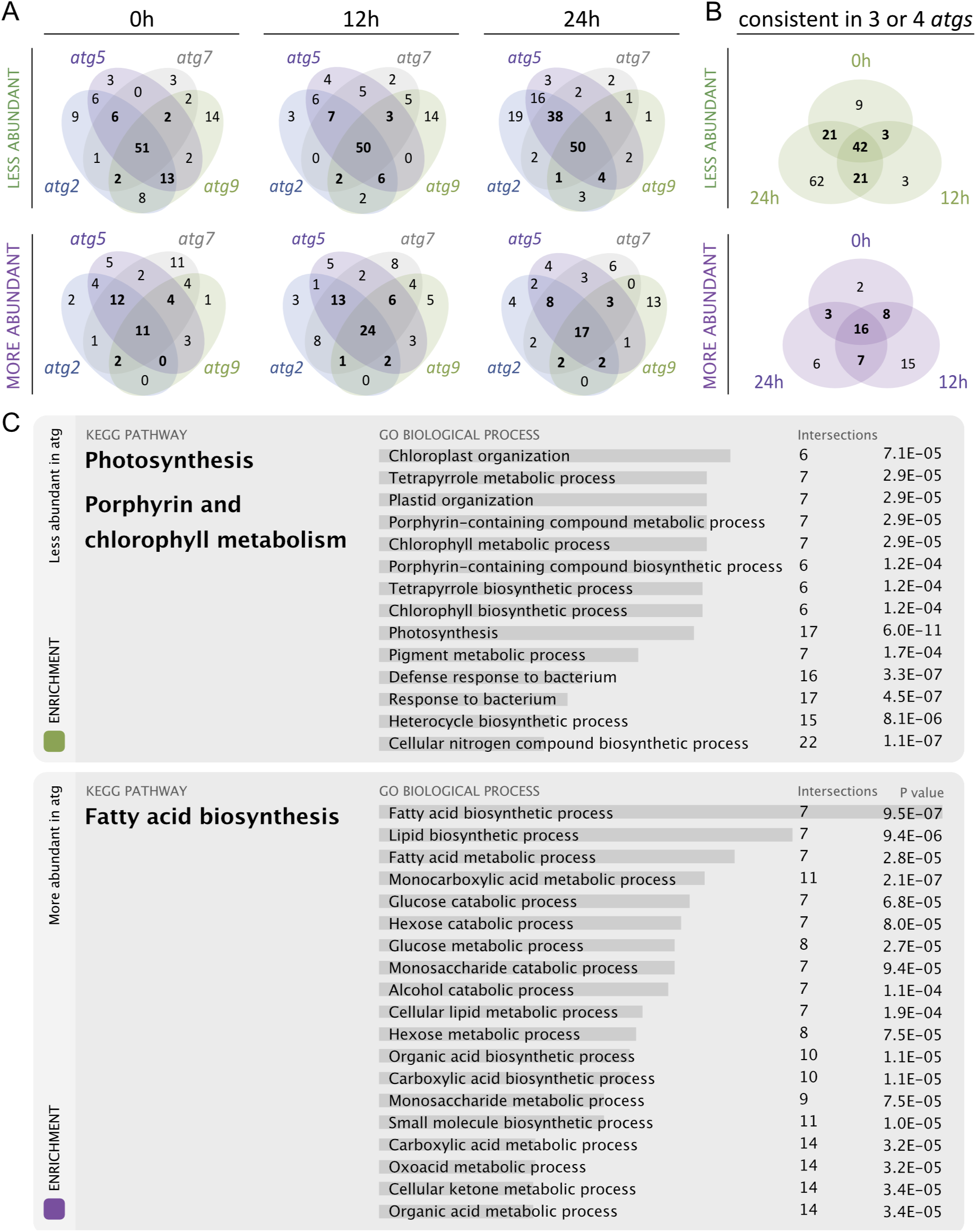
Differentially enriched plastid proteins in autophagy mutants during the dark-light transition. A. Venn diagrams for DEPPs at 0, 12 and 24h for the individual autophagy mutants. B. Venn diagrams for DEPPs common to at least three atg lines at the indicated times. C. Enrichment of metabolic pathways and biological processes for the DEPPs of autophagy mutants that were consistent in at least two time points. The horizontal bars behind the text indicate the Enrichment score for each BP.

We constructed a heatmap representing Z-scores of each family group of these DEPP set that were consistent over time and *atg* lines. The heatmap clearly showed two clusters, one of them including all the less abundant plastid proteins and the other one containing the more abundant plastid proteins (**Figure 5**). From the heatmap it is clear that the greatest differences in terms of less abundant plastid proteins are seen at 0h, and that *atg9* is in general the least different to WT (**Figure 5**). The less abundant plastid proteins consisted of 86 proteins including 28 proteins related to photosynthesis;12 proteins were related to the light phase of photosynthesis; 7 proteins were related to ATP synthesis; 6 were related to carbon fixation; 14 ribosomal proteins; 6 proteins involved in porphyrin and chlorophyll metabolism; and 6 proteins involved in amino acid metabolism (**Figure 5**).

**Figure 5.**
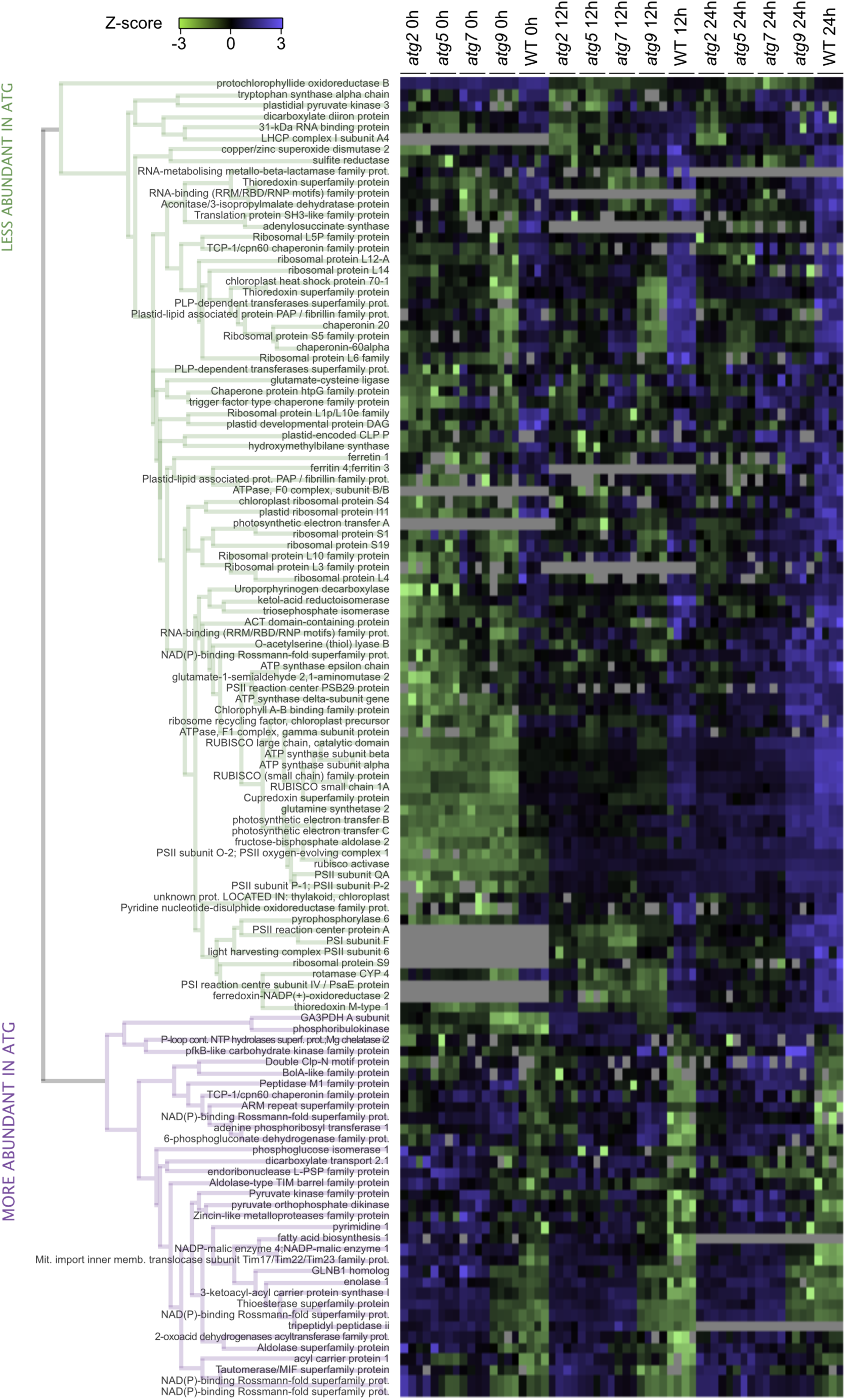
Heatmap and list of differentially enriched plastid proteins in autophagy mutants during the dark-light transition. The colours indicate the Z-score, which in some cases reached values much lower than -3 (e.g. -7) but were set from -3 to +3 to be more comprehensive. Grey colour indicates missing data. The hierarchical clustering generated two clear clusters which were separated in the figure as less and more abundant for visual purpose only.

The more abundant proteins were dominated by 8 proteins involved in carboxylic acid metabolism and biosynthesis, 8 proteins involved in carbohydrate metabolism and catabolism, 2 related to carbon fixation, and 2 involved in amine biosynthesis (**Figure 5**). Altogether, the lower abundance of proteins involved in photosynthesis and chlorophyll biosynthesis confirms the delayed chloroplast development in *atg* lines that has been suggested by the physiological data and the lower number and smaller chloroplast (**Figure 2E**) and the lower photosynthetic activity (**Figure 3**). The higher abundance sets of proteins indicate that the effect on altered chloroplast development in *atg* lines affect biochemical pathways differentially and do not simply imply less plastids or plastid size.

### The proteomic data is not explained by transcriptional regulation

Given the high number of plastid proteins less abundant in autophagy mutants, we decided to investigate matched transcriptomics of these plants to see if there is an explanation for the DEPP set through altered transcriptional regulation. Therefore, we analysed transcriptomic data (**Supp. File 4**) for the subset of DEPP defined for **Figure 5** and compared this data with the equivalent proteomics data. However, there was no overall correlation between DEPPs and their transcripts (**Figure 6A**). Nevertheless, it was clear that transcripts for less abundant plastid proteins were typically more abundant at 12h in *atg* lines than in WT, indicating that lower protein abundances were not directly due to transcriptional downregulation in *atg* lines.

**Figure 6.**
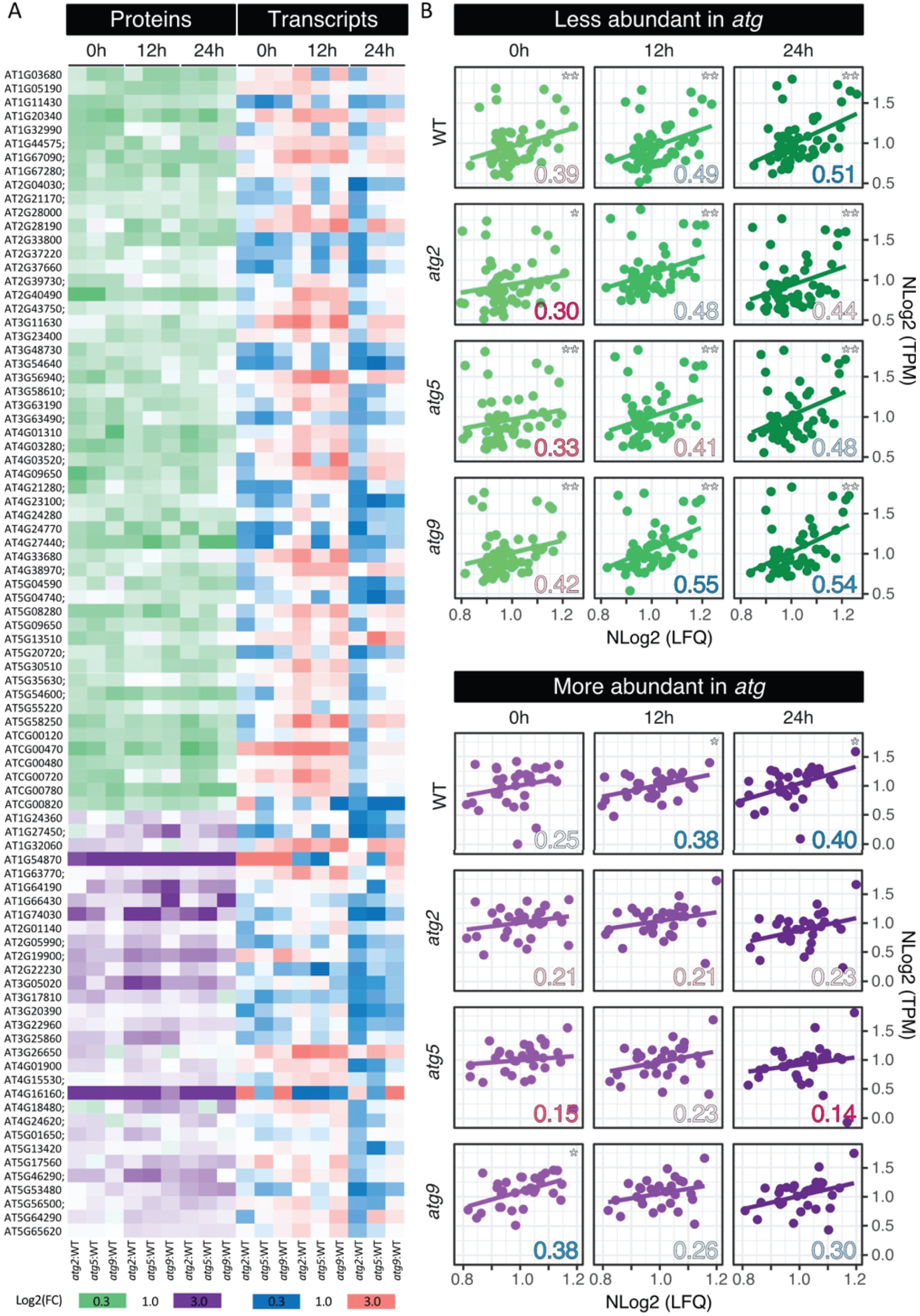
Correlation between proteomic data and transcriptomic data for differentially enriched plastid proteins. **A.** Heatmap showing the Log2(FC) for the proteomic and transcriptomic data of each autophagy mutant relative to WT at 0, 12 and 12h. **B.** Correlation graphs expressing transcriptomic vs proteomic data as normalized Log2(TPM) vs normalized Log2(LFQ) for the DEPP in WT and *atg* lines at 0, 12 and 24h. The numbers in the right bottom of each graph indicates the Pearson correlation coefficient and the colours indicate lowest (red) and greatest (blue) values. The asterisks in the top right corner indicate the p- value as follows *, p-value < 0.05 and **, p-value < 0.01.

To further explore correlations, we used normalized proteomic and transcriptomic data and performed Pearson correlation tests and evaluated their significance (**Figure 6B**). For the set of less abundant plastid proteins in *atg* lines, there was a Pearson correlation between the proteins and transcripts in all the cases (at a *p*-val=0.05, **Figure 6B**) but this correlation was often better for WT than for *atg* lines. For all the lines, the Pearson correlation was worse at 0h than at 12h and 24h (**Figure 6B**). Also, for this feature, *atg9* was the one line that was most similar to WT (**Figure 6B**). The correlation for the more abundant proteins was lower than that for less abundant proteins (**Figure 6B**). In particular, *atg2* and *atg5* never showed a significant correlation between protein and transcript abundances and only *atg9* showed a significant correlation at 0h (*p*-val=0.05) but not at 12h and 24h (**Figure 6B**). The WT showed a significant correlation at 12 and 24h (*p*-val=0.05) but not at 0h (**Figure 6B**). At *p*- value of 0.01, none of the lines showed a significant correlation.

Overall, the transcriptomic data highlights the importance of analysing proteomic data in autophagy mutants to gain understanding of the molecular pathways that are differentially expressed, as previously noted in maize autophagy mutant analysis (McLoughlin et al. 2018, 2020). In general, the results suggest that the transcriptional photomorphogenesis programs for plastid-localised gene products continues in autophagy mutants, and their proteomic differences are generally not explained by a differential transcription of the associated genes. To us, two main factors could be affecting the photomorphogenic process in *atg* lines, one could be the disruption of lipid membrane machinery that makes thylakoids ready to accept proteins, and the other could be an energetic restriction in *atg* lines that limits the translation/assembly of photosynthetic protein machinery.

### Analysis of de-etiolation phenotype in lipid metabolism single mutants sdp1 and tgd1, and lipid metabolism and autophagy double mutants

Autophagy mutants are known to have less triglyceride (TAG) and reduced TAG mobilization (Fan et al 2019). Thus, we tested if a mutant for a triacylglycerol lipase, *sugar dependent 1* (*sdp1*), not having less TAG but having impaired the capacity of using them (i.e, reduced mobility), could mimic the greening phenotype of autophagy mutants. On the other hand, if chloroplast biogenesis is slowed in autophagy mutants because they lack lipids to build thylakoids, mutants for the chloroplast lipid permease *trigalactosyldiacylglycerol1* (*tgd1*) might mimic the phenotype of *atg* lines, and *tgd1/atg2 and tgd1/atg5* lines should further exacerbate the phenotype. Therefore, we also tested the phenotype of available *tgd1* single mutants and *tgd1atg2* and *tgd1atg5* double mutants (Fan et al 2019) during the dark to light transition to test these ideas.

The lipid metabolism mutant *sdp1* and double mutants *sdp1/atg2* and *sdp1/atg5* were affected in their ability to green after transfer from dark to light, even more than autophagy mutants (**Figure 7A, Supp. Table 1**). The lipid mutant *tgd1* and the double mutants *tgd1/atg2 and tgd1/atg5* also showed lower GI compared to WT, but were not as affected as *sdp1* mutants (**Figure 7A, Supp. Table 1**). In terms of the photosynthesis over respiration ratio, we observed that at 12h *sdp1* and double mutants *sdp1/atg2* and *sdp1/atg5* had lower values than WT plants and in some cases even lower values than autophagy mutants (**Figure 7B**). At 24h the differences persisted for the double mutants *sdp1/atg2* and *spd1/atg5* (**Figure 7B**). No differences were observed for *tgd1* single mutant at both 12 and 24h, and although the ratio was slightly lower for the double mutants *tgd1/atg2* and *tgd1/atg5*, statistical differences were only found at 12h for *tgd1/atg5*. Similar results were obtained in terms of maximum quantum yield of PSII. At 6 and 12 h, all the single mutants *atg2*, *atg5*, *sdp1* and the double mutants *spd1/atg2 spd1atg5* had lower F_V_/F_M_ values than WT, and the differences persisted for the double mutants at 24h (**Figure 7C**). However, the F_V_/F_M_ values for *tgd1* were similar to WT at all times and only at 12h were differences observed between the double mutants *tgd1/atg2* and *tgd1/atg5* (**Figure 7C**). Likewise, ETR was generally lower in autophagy mutants, *sdp1* and *sdp1atg2* and *sdp1atg5* double mutants (**Figure 7D**). However, *tgd1*, *tgd1/atg2 and tgd1/atg5* mutants showed ETR values more similar to WT (**Figure 7D**).

**Figure 7.**
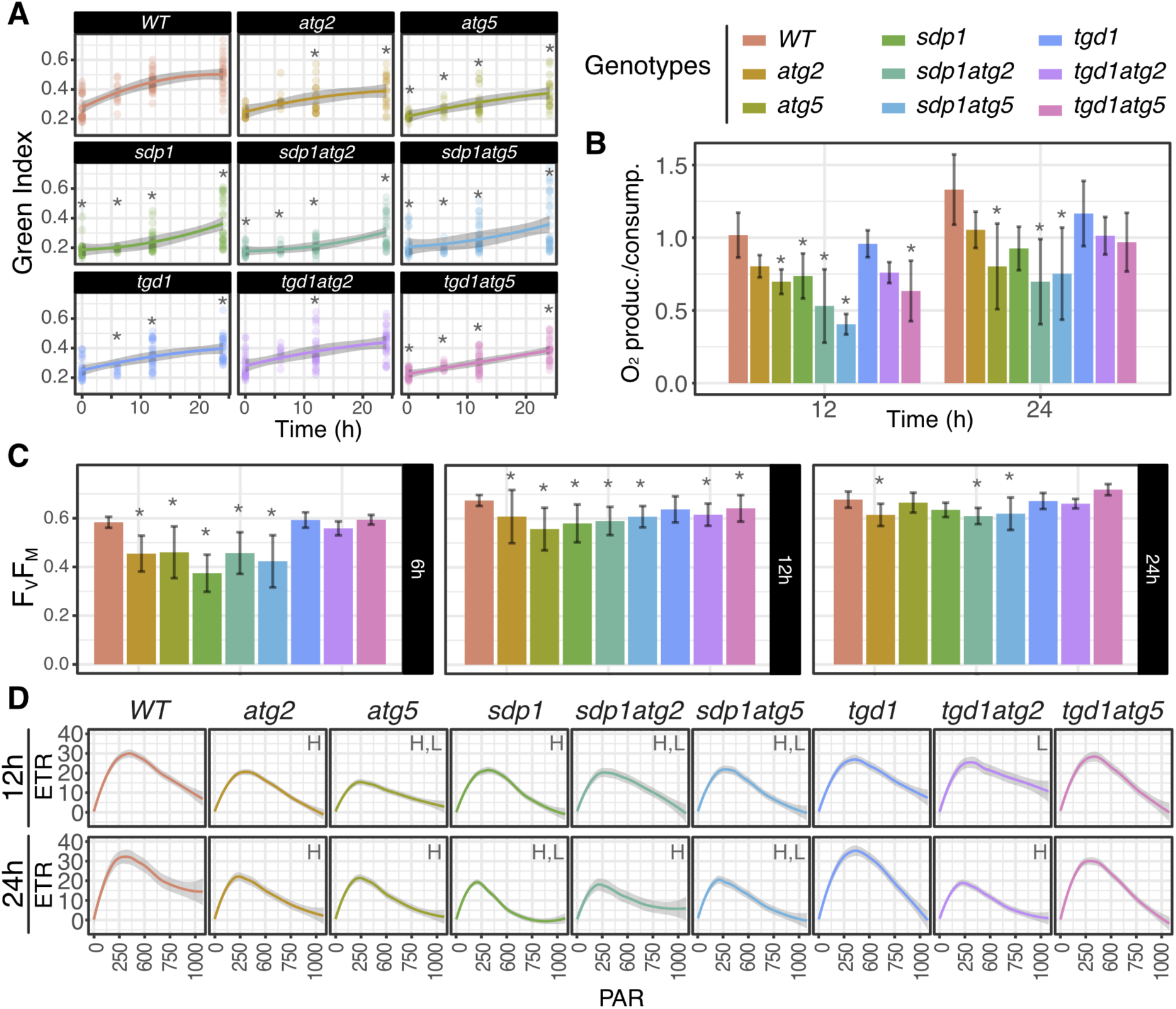
**A.** Green Index for WT, autophagy and lipid mutants, and autophagy/lipid double mutants at 0h, 6h, 12h and 24h of light treatment. **B.** Ratio between photosynthetic activity and respiration for WT, autophagy and lipid mutants, and autophagy/lipid double mutants at 12h and 24h. **C.** Analysis of F_V_/F_M_ values for WT, autophagy and lipid mutants, and autophagy/lipid double mutants at 6h, 12h and 24h. **D.** Relative apparent electron transport rate (ETR) plots for WT, autophagy and lipid mutants, and autophagy/lipid double mutants at 12h and 24h. The grey regression represents +/- 99% confidence level intervals for prediction for the polynomial regression and H, and L, represent respectively significantly higher values for *WT* respect to autophagy mutants in terms of the height (H) and length (L) of the maximum ETR as explained in Figure S3 and S5. Asterisks indicate statistically significant differences (P < 0.05) by Tukey’s HSD test compared to WT.

Together, these results show that *tgd1* mutants mimick the greening phenotype of autophagy mutants but are not affected in terms of photosynthetic capacity, whereas the double mutants display very similar phenotypes to the *atg* single mutants (i.e. *atg2*∼ *tgd1/atg2*, and *atg5*∼*tgd1/atg5*, **Figure 7A-C, Supp. Table 1**). Thus, for most of the parameters the addition of mutations in *atg* genes to *tgd1* mutants negatively affected its phenotype and almost mimicked its respective *atg* mutant (**Figure 7B-C**). This suggests that the *atg* mutations have more impact than the *tgd1* mutation in photomorphogenesis. In contrast, *sdp1* mutants also displayed a delayed greening phenotype but it was clearly more pronounced than in the autophagy mutants (**Figure 7, Table S1**). The addition of mutations in *atg* genes to *sdp1* mutant did not rescue the phenotype, instead the phenotype was mostly similar to *sdp1* single mutants and worse than *atg* single mutants (**Figure 7A, Supp. Table 1**). This suggests that impaired mobility of TAG has a stronger effect on photomorphogenesis than the lack of autophagic activity, and also suggests that the reduced TAG mobility of *atg* mutants may contribute to their delay in photomorphogenesis.

### Lowered carbon availability exacerbates the delayed greening effect in autophagy mutants

Given that our study on *sdp1* mutants suggested that the delay in photomorphogenesis of *atg* lines may be related to their reduced capacity to degrade TAG, we tested the greening phenotype under different levels of sucrose to see if lower levels of sucrose further compromised their phenotype and/or if higher levels of sucrose can attenuate their phenotype. We analysed the greening of WT and autophagy mutants during the dark to light transition (0, 6, 12 and 24h post illumination) under normal conditions (0.5% Suc), poor carbon source (C) conditions (0.1% Suc) and rich C conditions (2% Suc). At 0.5% Suc, the greening of the *atg* lines is delayed but all of them reach WT values by 24h (**Figure 8A**). When subjected to the poor C medium the differences between WT and autophagy mutants were exacerbated, all of them being statistically different at 6 and 12h and even one of them at 24 h (*atg9*), whereas when subjected to a richer C medium the differences with WT were completely lost (**Figure 8A**). Likewise, the lower F_V_/F_M_ of autophagy mutants observed at 6h and 12h in 0.5% sucrose are also observed in 0.1% sucrose but not in 2% sucrose (**Figure 8B**). As observed before, the differences in F_V_/F_M_ were reduced over the time becoming insignificant by 24h except for 0.1% sucrose where some differences between WT and autophagy mutants persist (**Figure 8B**). The ETR was also reduced in autophagic mutants compared to WT plants, with the differences more significant at earlier times post illumination and at lower sucrose level (**Figure 8C**).

**Figure 8.**
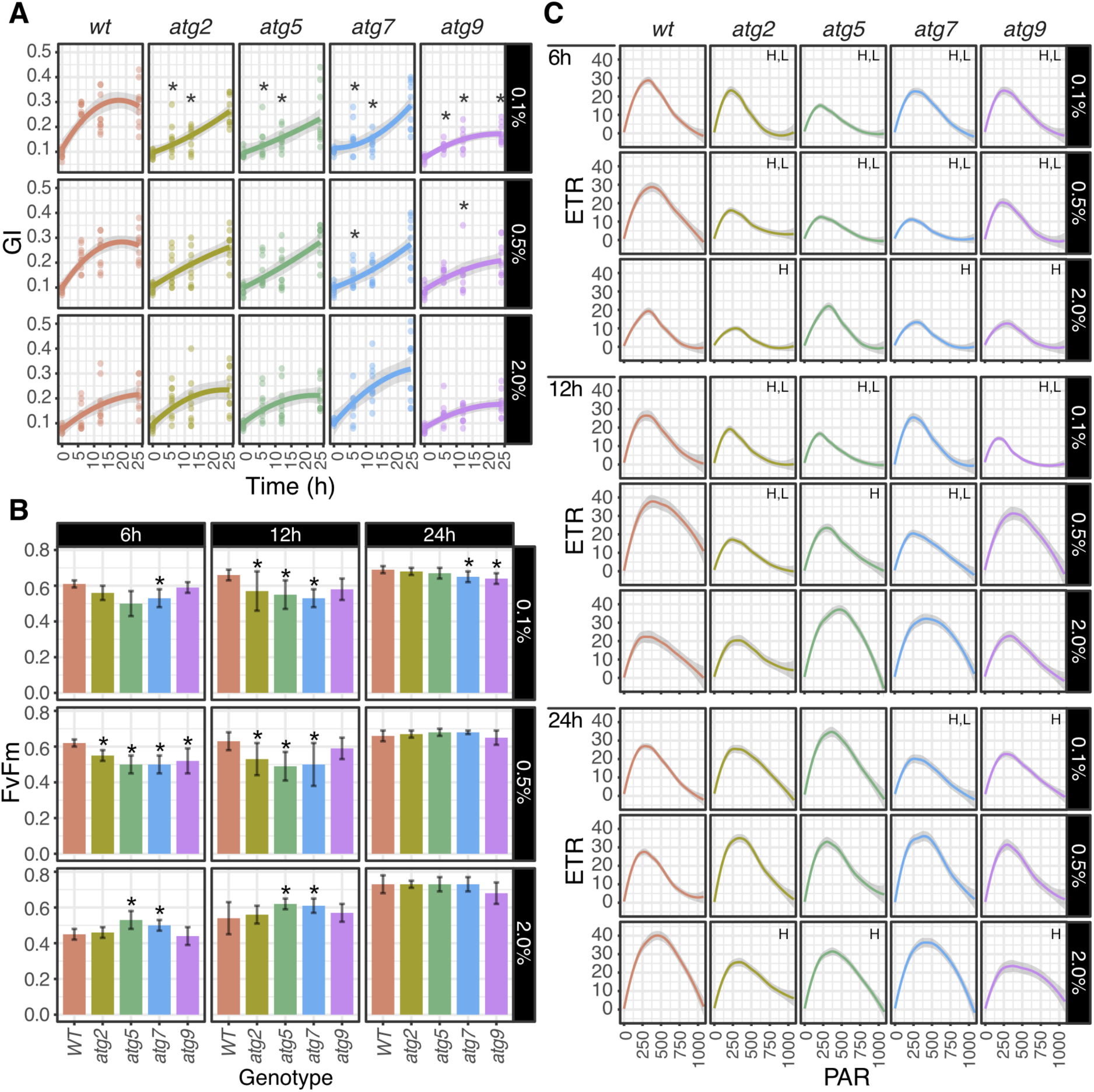
High carbon source (C) medium suppresses the delay in photomorphogenesis observed in autophagy mutants, whereas low C medium worsens the atg phenotype. **A.** Green Index under different C nutrient regimes. **B.** Analysis of F_V_/F_M_ under different C nutrient regimes. **C.** Relative apparent electron transport rate (ETR) plots for WT and autophagy mutants. In all the cases the genotypes were evaluated at 0h, 6h, 12h and 24h post illumination in mediums containing 0.1%, 0.5% and 2.0% sucrose. Asterisks indicate statistically significant differences (P < 0.05) by Tukey’s HSD test compared to WT. For ETR, the grey regression represents +/- 95% confidence level intervals for prediction for the local regression and H, and L, represent respectively significantly higher values for *WT* respect to autophagy mutants in terms of the height (H) and length (L) of the maximum ETR as explained in Figure S3 and S6.

Altogether, the data show that a high level of sucrose can ameliorate the delay in greening observed in autophagy mutants confirming that C restrictions in autophagy mutants are one of the reasons why autophagy mutants display a delayed photomorphogenesis.

## Discussion

### Autophagy mutants have delayed chloroplast development during photomorphogenesis

While the maturation from etioplast into chloroplast in cotyledons of seedlings during the first hours after light exposure is a well-documented process (Rudowska et al. 2012; Pogson et al. 2015), a role of autophagy in this process remains largely unexplored. When we found that plastid organelle markers abundance was the most affected during the dark to light transition of young autophagy mutant seedlings (**Figure 1**, **Figure S1** and **Figure S2**), we sought to further explore the role of autophagy in photomorphogenesis with a special focus on chloroplast development.

Chloroplasts house the machinery of photosynthesis as well as nitrogen assimilation, fatty acid and amino acid biosynthesis, among other process, thus their development is known to strongly influence plant development (Pogson et al. 2015). Chloroplast development requires multiple molecular processes such as nuclear gene expression, protein import, plastid gene expression, chloroplast RNA processing and ribosome assembly, protein maturation and degradation, thylakoid biogenesis, lipid biosynthesis, chlorophyll biosynthesis, metabolite transport, and photosystem assembly (Waters and Langdale 2009). Here, we observed that autophagy mutants had lower chlorophyll content, a reduced number and size of chloroplasts (**Figure 2**) and poorer light-phase dependent photosynthetic activity during photomorphogenesis (**Figure 3**), suggesting that one or more of these molecular processes involved in chloroplast development was affected in autophagy mutants. The lower PSII maximal efficiency in autophagy mutants, suggest either that the PSII is damaged or not fully developed at 12h in these lines (**Figure 3B**), potentially due to lower chlorophyll levels in autophagy mutants at 12h (**Figure 2D**). Moreover, at 12h, the higher **φ**NO results observed in autophagy mutants revealed that the photochemical energy conversion and protective regulatory mechanism in these mutants were less efficient than in the WT either due to a less active PSII reaction centre or a poorer trans-thylakoidal proton gradient (**Figure 3B**). As a consequence, the electron transport rate and photosynthetic activity in these mutants were lower than in WT (**Figure 3C and D**). Our data suggest that while autophagy mutants can complete photomorphogenesis, it is delayed compared to WT. From the literature it is known that by 3 to 5 weeks old, chlorophyll content is no longer reduced and there are no changes in the transcripts for chlorophyll biosynthesis and degradation in *atg2* (Jiang et al. 2020).

### Autophagy mutants show lower abundance of chlorophyll biosynthetic and photosynthetic proteins and increased abundance of proteins involved in plastid fatty acid biosynthesis

As autophagy acts post-transcriptionally to alter the protein complement of cells, it was important to study the abundance of plastid proteins to further understand the delayed greening identified in autophagy mutants. We observed that these mutants have lower abundance of proteins involved in chlorophyll biosynthesis and photosynthesis but greater abundance of proteins involved in fatty acid biosynthesis and carbohydrate metabolism (**Figure 4** and **Figure 5**). These findings are in agreement with previous reports using adult maize autophagy mutants that showed reduced levels of thylakoidal proteins and increased levels of proteins involved in fatty acid metabolism (McLoughlin et al. 2018). Basal autophagy has been previously shown to contribute to TAG synthesis (Fan et al. 2019), thus it is possible that the observed increase in fatty acid biosynthetic proteins is a response to compensate for the lack of autophagy-mediated TAG synthesis. On the other hand, we observed that the differential abundance of proteins in autophagy mutants is not simply explained by transcriptional regulation (**Figure 6**). In general, the correlation at 0h between protein and transcript abundance were poor for all genotypes (**Figure 6B**). At 12h and 24h after the transfer to light, the correlation for the less abundant proteins improved compared to 0h. However for the more abundant proteins, only WT showed a considerable improvement of the correlation between protein and transcript abundance at 12h and 24h compared to 0h, suggesting that some of these proteins may be degraded by autophagy in the WT plants but not in *atg* lines influencing the poorer correlation. Hence, it is unlikely that the differential expression of photosynthetic related proteins is related to disrupted signalling events in autophagy mutants resulting in lower transcription. Instead, it is more likely that the altered lipid metabolism in autophagy mutants affects the thylakoid structure and limits the assembly of photosynthetic complexes. A link between autophagy, photosynthesis and fatty acid has been established in the model alga *Chlamydomonas reinhardtii*, in which the inhibition of fatty acid biosynthesis was shown to downregulate photosynthesis and induce autophagy (Heredia-Martínez et al. 2018). In particular, the authors concluded that a change in the monogalactosyldiacyl-glycerols / digalactosyldiacyl-glycerols ratio (MGDG/DGDG) was responsible for altering the thylakoid structure and thus the photosynthetic activity (Heredia-Martínez et al. 2018). As autophagy is known to affect fatty acid content and composition in *Arabidopsis* (Minina et al. 2018; Fan et al. 2019; Havé et al. 2019), it is possible that the differing lipid profiles in autophagy mutants are partially responsible for the delayed chloroplast development and the resulting lower photosynthetic activity as observed in *C. reinhardtii*.

### The delayed photomorphogenesis in autophagy mutants is probably not related to limitations in lipid import into the chloroplast or lower MGDG and DGDG content

TGD1 is a chloroplastic permease (inner envelope) involved in the unidirectional lipid transfer from the ER to plastids, necessary for thylakoids formation (Xu et al. 2003, 2010). The TGD complex is known to transport phosphatidylcholine (PC) but it is expected that other MGDG precursors are also transported (Michaud and Jouhet 2019). Potential candidates are phosphatidates (PA) and diacylglycerols (DAG) (LaBrant et al. 2018). These precursors are used to produce MGDG and DGDG which together compose 80% of chloroplast lipid and are essential for the light phase of photosynthesis (Li and Yu 2018) (**Figure 9**).

**Figure 9.**
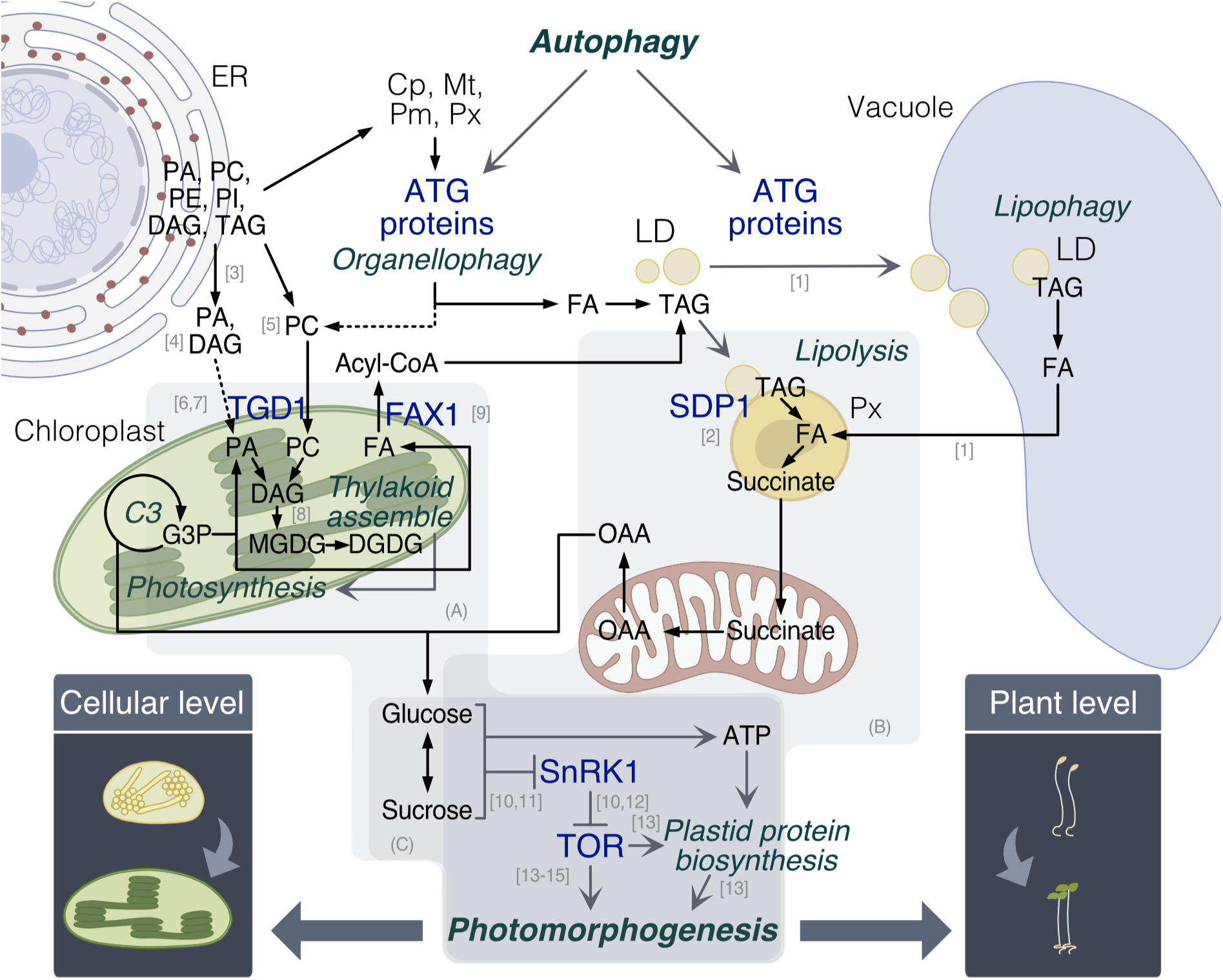
Mechanistic insights into the autophagy effect on photomorphogenesis and its relation with the current literature. The scheme describes the main mechanism discussed here by which autophagy could contribute to photomorphogenesis. The first mechanism is the contribution of autophagy to phosphatidylcholine recycling via organellophagy, which is used for thylakoid assembly during chloroplast development. We showed that *tgd1* lines presented a delayed greening (Fig. 7) supporting the mechanism enclosed in (A). The second mechanism involves the contribution of autophagy to fatty acids release, either through organellophagy or lipophagy. Both mechanisms converge in the production of sugars, either through photosynthesis or FA degradation, which in turn are known to support photomorphogenesis both energetically and as signalling molecules allowing TOR activity. Through the *sdp1* lines, we showed that lipolysis is indeed relevant for photomorphogenesis (Fig. 7) supporting the mechanism enclosed in (B). Finally, we showed that sugar levels can affect the degree of photomorphogenesis (Fig. 8) as shown in (C). [1]: Fan *et al*. 2019; [2]: Eastmond 2006; [3]: Li-Beisson *et al*. 2013; [4]: Michaud *et al*. 2019; [5]: LeBrant *et al*. 2018; [6]: Xu *et al*. 2003; [7]: Xu *et al*. 2010; [8]: Li *et al*. 2018; [9]: Schnurr *et al*. 2002; [10]: Baena-Gonzalez *et al*. 2007; [11]: Ghillebert *et al*. 2011; [12]: Nukarinen *et al*. 2016; [13]: Chen *et al*. 2018; [14]: Cai *et al*. 2017; [15]: Pfeiffer *et al*. 2016. Cp: chloroplasts; ER: endoplasmic reticulum; DAG: diacylglycerol; DGDG: digalactosyldiacylglycerol; FA: fatty acids; LD: lipid droplets; MGDG: monogalactosyldiacylglycerol; Mt: mitochondria; PA: phosphatidate; PC: phosphatidylcholine; PE: phosphatidylethanolamine; PI: phosphatidylinositol; Pm: plasma membrane; Px: peroxisome; SnRK1: Snf1-related kinase 1; TAG: triacylglycerols; TOR: target of rapamycin.

Because extensive lipid trafficking is required for thylakoid lipid biosynthesis (Awai et al. 2006) and the ER lipid assembly pathway was shown to be more relevant than the chloroplast pathway in the fatty acid biosynthesis and membrane lipid turnover under basal autophagy (Fan et al. 2019), we tested if *tgd1* mutants mimic the phenotype of autophagy mutants. It is worth noting that mature *tgd1* mutants do not show alterations in their photosynthetic parameters when compared to WT plants (Li et al. 2012). While gross carbon assimilation were shown to be marginally affected (10-15% less), Rubisco activity, electron transport rate and photosynthesis I and II oxidation status were not altered (Li et al. 2012). Hence, reduced photosynthetic capacity in *tgd1* during de-etiolation would suggest a developmental delay. Our results showed that *tgd1* mutants mimicked the slow greening phenotype of autophagy mutants (**Figure 7A**). Moreover, the *tgd1* mutant is known to display a upregulation of the chloroplastic lipid biosynthetic pathway (Xu et al. 2003), causing an accumulation of TAG (Xu et al. 2005). We also observed a greater abundance of proteins involved in chloroplastic fatty acid biosynthesis (**Figure 4** and **Figure 5**). Thus, the similar phenotype of *tgd1* and autophagy mutants as well as the induction of the chloroplastic lipid biosynthetic pathway in these mutants point to a possible limitation in the lipid trafficking between the ER and the chloroplast in autophagy mutants that may delay chloroplast development. However, we would note that photosynthetic performance was not affected in *tgd1* mutants (**Figure 7B-D**). Thus, we conclude that limitations in neither chloroplast lipid import nor MGDG-DGDG content are solely responsible for the developmental delay in autophagy mutants compared to WT.

### Lipid turnover plays a role in photomorphogenesis and the contribution of autophagy to lipid homeostasis may be the reason of its role in photomorphogenesis

Autophagy mutants are known to have less TAG and reduced TAG mobilization (Fan et al 2019). We observed that autophagy mutants maintained higher abundance of proteins involved in plastid fatty acid biosynthesis (**Figure 4** and **Figure 5**), which could be indicative of a compensatory response to lower levels of fatty acids. The increase of transcripts for lipid biosynthetic enzymes was also recently observed in autophagy mutants (*atg12*) of maize subjected to carbon starvation (McLoughlin et al. 2020). Based on the hypothesis that the different levels and profiles of lipids may affect photomorphogenesis process in autophagy mutants, we assessed greening in the *sdp1* mutants, which like autophagy mutants have impaired capacity to use TAG, to see if they mimic the slow greening phenotype of autophagy mutants. Our results showed that *sdp1* mutants also have impaired photomorphogenesis (**Figure 7**), so it is possible that the autophagy-related slow-greening phenotype we observed is due to a limitation in TAG catabolism (**Figure 9**). In fact, the *sdp1* phenotype was more severe than the one observed in autophagy mutants. *Sdp1* encodes for a lipase that controls fat storage breakdown (Eastmond 2006), and the *sdp1* mutant displays an 80% reduction in lipid hydrolysis rates compared to WT (Li-Beisson et al. 2013). Although we know autophagy mutants have lower TAG content (Minina et al. 2018; Fan et al. 2019), it is not expected that the autophagy mutants would have such a strong reduction in TAG catabolism as *sdp1.* Therefore the more severe slow greening phenotype in *sdp1* than in autophagy mutants might be expected if TAG availability is the explanation of the phenotype.

### Energetic restrictions in autophagy mutants also contribute to the delayed photomorphogenesis

Our results showed that the limitations in TAG catabolism (*spd1*) have more severe consequences in photomorphogenesis than the limitations in lipid trafficking (*tgd1*). SDP1 is the major TAG lipase involved in lipid reserve mobilization during seedling establishment (Eastmond 2006; Thazar-Poulot et al. 2015). The *sdp1* mutants were isolated as mutants displaying a postgerminative growth arrest when sugars are not provided in the medium (Eastmond 2006). This is because SPD1 is ultimately responsible for sucrose production during seedling establishment. In particular, SDP1 degrades TAG to release fatty acids which are transferred to the peroxisome to be converted into succinate, which is used for the synthesis of carbohydrates (**Figure 9**). Moreover, the physical interaction between lipid droplets and peroxisomes, and thus SDP1 activity, is tightly regulated by sucrose levels, by way of high sucrose levels reducing this interaction, mediated by actin filaments (Cui et al. 2016). Therefore, there is significant evidence that SDP1 plays a role in controlling the energetic status of seedlings and now we can infer that this energetic status is also important for optimal photomorphogenesis.

Since our seedlings were grown in the dark for 5 days, it is also possible that induced autophagy was playing a role in our experimental conditions to further support the energetic requirement for the photomorphogenic process. In case of carbon deprivation, lipid droplets (or oil bodies) are degraded into fatty acids to produce energy through their catabolism and a specific type of induced autophagy, macrolipophagy, was shown to mediate this degradation (Fan et al. 2019). In this case, the lipid droplets are degraded in the vacuole rather than in the peroxisomes (**Figure 9**). Thus, it would be expected that SPD1 metabolism is induced concomitantly with induced-autophagy and both pathways contribute cellular resources and energy to sustain photomorphogenesis. To test if energetic restriction in autophagy mutants contributes to the delay in photomorphogenesis we repeated the characterization of autophagy mutants under lower and higher levels of C. Our results confirmed that lower levels of C exacerbate the greening phenotype of autophagy mutants whereas much higher levels of sucrose can ameliorate them (**Figure 8**). Therefore, it is possible that the lower capacity of *atg* seedlings to produce sugars through lipophagy, results in lower capacity to produce ATP to sustain protein biosynthesis (**Figure 9**).

Very recently, mitophagy was shown to take place in cotyledons during de-etiolation (6h post illumination) and an analogous delay in greening capacity of *atg5* was presented (Ma et al. 2021), suggesting that autophagy plays a role in adjusting the number of mitochondria during photomorphogenesis and proposing cross-talk with delayed plastid biogenesis. We show here that the *atg* greening phenotype is not specific to *atg5* and is not constitutive, but conditional on C status. In addition, we observed that respiration rate per se is not affected in *atg* mutants compared to WT during de-etiolation at least at 12 hours (Figure S4), suggesting that the possible changes in mitochondrial turnover do not affect the overall respiration rate and that mitophagy is unlikely to be the primary process determining the slow greening phenotype. However, the phenotype is likely to relate to the function of both organelles in the context of sugar availability and energetic supply to biosynthetic processes.

Sugars not only play an energetic role but also act as signalling molecule in plants (Baena-González et al. 2007). For instance, sucrose promotes the accumulation of trehalose-6- phosphate (T6P) which inhibits Snf1-related kinase 1 (SnRK1), an antagonist of target of rapamycin (TOR), capable of repressing biosynthetic processes and plant growth (Baena-González et al. 2007; Ghillebert et al. 2011). On the other hand, TOR complex promotes metabolic activity and growth, ribosome biogenesis, protein synthesis, and influence the rate of respiration with different substrates (Lastdrager et al. 2014, O’Leary et al 2020). In the first hours of Arabidopsis seedlings transitioning from dark to light, TOR has been shown to be responsible for the phosphorylation of ribosomal protein S6 (RPS6) to enhance protein translation to establish the photosynthetic apparatus (Chen et al. 2018). Therefore, it is possible that the suppression of lipophagy in *atg* lines, induces a starvation-like condition in *atg* etiolated seedlings, that induces SnRK1 activity, which in turns phosphorylate and inactivate TOR reducing the translation of the photosynthetic machinery (**Figure 9**). This putative process would implicate an important role of autophagy and lipophagy in seedling establishment and deserves further exploration.

## Conclusion

We conclude that autophagy influences the rate of etioplast to chloroplast conversion and plays a role in photomorphogenesis most probably by contributing to lipid mobilization and energy production (**Figure 9**). The effect could be both; indirectly through the modulation of cellular processes that contribute to cellular resources for growth and development, and also directly by etioplasts maintaining higher levels of lipid synthesis machinery to provide TAG for thylakoid membranes and limiting the availability of reductant and ATP inside the plastid for use in the biosynthetic demands of protein import, protein synthesis and photosynthetic protein complex assembly.

## Materials and Methods

### Plant materials and growth conditions

We used arabidopsis (*Arabidopsis thaliana*) seeds of Col-0 as WT, four autophagic mutants *atg2-1* (SALK_076727, Yoshimoto et al. 2009), *atg5-1* (SAIL_129B07, Chen et al. 2015), *atg7-2* (GABI_655B06, Lai et al. 2011) and *atg9-2* (SALK_130796, Zhuang et al. 2017) which we referred as *atg2*, *atg5*, *atg7* and *atg9* respectively, a lipid thylakoid permease mutant *tgd1* (Xu et al. 2003), a TAG lipase mutant (*sugar-dependent 1-4*) *sdp1-4* (Eastmond 2006; Fan et al. 2017) and the respective lipid and autophagy double mutants *tgd1atg2*-1, *tgd1* atg5-1, sdp1atg2-1, and sdp1 atg5-1 (Fan et al. 2019).

Growth conditions vary between the developmental transition studied. Three developmental transitions were established; during early seedling development (T1), dark to light greening of seedlings (T2) and light to dark senescence of leaves (T3). For T1 and T2, seeds were surface-sterilized and carefully dispensed on a stainless-steel wire mesh platform (mesh size 1 mm; 3 cm x 3 cm x 3 cm) layered previously with 1% sterilized agarose in a round plastic vessel containing 300 mL of liquid medium (1⁄4- strength Murashige and Skoog -MS- medium without vitamins, 1⁄4- strength Gamborg B5 vitamins solution, 2 mM MES, 1% [w/v] sucrose, pH 5.8). For T1, plants were grown under a 16/8 h light/dark period with a light intensity of 100-125 μmol·m^-2^·s^-1^ at 22°C and samples were collected two days after germination (T1_0d) and then one, three and five days after (T1_1d, T1_3d and T1_5d). For T2, seeds were grown five days in dark (T2_0d), and then grown under 16/8 h light/dark period with a light intensity of 100-125 μmol·m^-2^·s^-1^ at 22°C for one, three and five days (T2_1d, T2_3d, T2_5d).

For T3, plants were grown in Hoagland’s solution and after 21 days (T3_0d_UC), leaves were covered (leaf number three, five, and seven) (C) by aluminum foil to study dark-induced senescence. Covered leaves (leaf number: three, five, seven) and uncovered (UC) leaves (leaf number: four, six, eight) were collected over time course (1 day, 3 days and 5 days) (Uncovered leaves: T3_1d_UC, T3_3dU_C, T3_5d_UC, Covered leaves: T3_1d_C, T3_3d_C, T3_5d_C).

For successive physiological study of the autophagy mutants and all the study of the lipid metabolism mutants, the growth conditions were the same but they were sown on plates containing 0.5X MS medium supplemented with 0.5% sucrose (unless otherwise stated) and 0.8% plant agar. For this, seeds were surface sterilized by 1 min of 70% ethanol, rinsed once with sterile water, 3 min with 20% sodium hypochlorite, and washed five times with sterile water.

### Protein isolation and sample preparation

Approximately 200 mg of frozen tissue were ground under liquid nitrogen before extraction in 400 μL of 125 mm Tris–HCl pH 7.5, 7% (w/v) SDS, 10% (v/v) β-mercaptoethanol, 0.5% (w/v) PVP40 with Roche protease inhibitor cocktail added at 1 tablet per 50 mL of extraction buffer. Protein extraction was carried out by chloroform/methanol extraction (Wessel and Flügge 1984) before washing the pellet twice in 80% (v/v) acetone.

Samples were resuspended in freshly prepared buffer (7M Urea, 2M Thiourea, 50mM NH_4_HCO_3_, and 10mM DTT) to obtain 15-25 μg/μL range after resuspension. Protein concentration was determined by Bradford assay and spectrophotometric measurement at a wavelength of 595 nm using bovine serum albumin as a standard.

200 μg of protein were treated with 25mM Iodoacetamide for 30 min in the dark. The sample solutions were then diluted to below 1 M urea with 50 mM NH_4_HCO_3_. For protein digestion, 10 mg of trypsin (dissolved in 0.01% [v/v] trifluoroacetic acid to a concentration of 1 mg/mL) was added to each sample and incubated at 37°C overnight. The samples were acidified to 1% (v/v) with formic acid and solid-phase extraction cleaned using Silica C18 Macrospin columns (The Nest Group). After each of the following steps, solid-phase extraction columns were centrifuged for 3 min at 150 g at room temperature. Before loading, the sample columns were washed with 750 μL of 70% (v/v) acetonitrile, 0.1% (v/v) formic acid and charged with 750 μL of 5% (v/v) acetonitrile, 0.1% (v/v) formic acid. After loading the samples onto the columns, two washes with 750 μL of 5% (v/v) acetonitrile, 0.1% (v/v) formic acid were carried out, followed by two elution steps with 750 μL of 70% (v/v) acetonitrile, 0.1% (v/v) formic acid. The eluate was dried under vacuum and resuspended in 5% (v/v) acetonitrile, 0.1% (v/v) formic acid to a final concentration of 1 μg/μL for MS.

### LC-MS/MS for dynamic MRM analysis

Using an Agilent 1290 Infinity II LC system, 10 μg of each sample were loaded onto an Agilent AdvanceBio Peptide Map column (2.1 x 250mm 2.7-Micron, P.N. 651750- 902), which was heated to 60 °C. Peptides were eluted over a 30 min gradient (0-15 min 3% [v/v] acetonitrile 0.1% [v/v] formic acid to 45% [v/v] acetonitrile 0.1% [v/v] formic acid; 15-15.5 min 45% [v/v] acetonitrile 0.1% [v/v] formic acid to 100% [v/v] acetonitrile 0.1% [v/v] formic acid; 15.5-16 min 100% [v/v] acetonitrile 0.1% [v/v] formic acid to 3% [v/v] acetonitrile 0.1% [v/v] formic acid; 16-30 min 3% [v/v] acetonitrile 0.1% [v/v] formic acid) directly into the Agilent 6495 Triple Quadrupole MS for detection.

### Analysis of targeted MRM data

Before conducting MRM analysis for the individual samples, a preliminary MRM analysis was performed using pooled samples (n=4) of all growth and developmental stages to obtain transition information, such as detectability in tissues and suitability of transition. For organelle-specific assay, 680 peptides (2231 transitions) representing 76 proteins were targeted. After obtaining the resulting MRM data, the final target list was selected using the following criteria: at least one peptide was selected per protein, and at least three transitions (one quantifier and two qualifiers) per peptide were chosen. Individual MRM analysis was performed using 221 transitions, corresponding to 31 proteins.

Raw data files from MRM analysis were processed using Skyline. Peak area integration was confirmed manually to correct potentially wrongly assigned targets. For further analysis, the integrated area of the quantifier ion was used. Normalization was performed to correct for differences between sample runs. Therefore, every peptide within a sample was divided by the median of all detected peptides in that sample. Then, to give peptides the same weight, normalized data were scaled by dividing each peptide by the median of that peptide across all samples.

### LC-MS/MS for shotgun proteomics

Digested peptides were resuspended in 20 μL of 2% acetonitrile (v/v), 0.1% formic acid (v/v), and 5 μl of this resuspension was loaded onto a C18 high capacity nano-LC chip (Agilent) in 98% Buffer A (0.1% (v/v) formic acid in Optima grade water (Fisher)) and 2% Buffer B (0.1% formic acid in Optima grade acetonitrile (Fisher)) using a 1200 series capillary pump (Agilent). Following loading, samples were eluted from the C18 column and into an inline 6550 Series QTOF mass spectrometer (Agilent) with a 1200 series nano pump (Agilent) using the following gradient: 2% B to 45% B in 26 minutes, 45% to 60% B in 3 minutes, 60% to 100% B in 1 minute. The samples were analyzed in sequence, and a blank was run between samples. The QTOF was operated in a data- dependent fashion with an MS spectrum collected prior to the three most abundant ions subjected to tandem mass spectrometry from doubly, triply, and higher charge states. Ions were dynamically excluded for 0.4 minutes following fragmentation. MS data were collected at eight spectra per second, while MS/MS spectra were collected at three spectra per second with a minimum threshold of 10,000 counts and a target of 25,000 ions per MS/MS event.

### Label-Free Quantitative shotgun proteomics data analysis

Raw data files were analyzed by the MaxQuant quantitative proteomics software package (Cox and Mann 2008). Peak lists were searched against the Arabidopsis Fasta database TAIR10_pep_20101214. Each genotype was represented by four biological replicates in three different times, 0h, 12h and 24h. Statistical analysis of Label-Free Quantitation (LFQ) values from MaxQuant was performed using the Perseus software (http://www.perseus-framework.org/) from MaxQuant (Tyanova et al. 2016). Proteins were filtered according to the presence in at least 70% of the samples tested and being annotated as plastid proteins according to SUBA4 (https://suba.live) consensus column. Subsequently, differential enrichment analysis between autophagy mutants and WT plants was performed and additional filtering was applied to retain those proteins that were differentially enriched in at least three out of four atg mutants and in at least two out of three times tested. Multiple-sample ANOVA test was used to determine significance. A protein was considered differentially expressed if the permutation-based false discovery rate (FDR) adjusted p-values were below 0.05. The resulting dataset was hierarchically clustered to produce heatmaps using the Z-score value in Perseus (Tyanova et al. 2016). Additionally, a GO term and KEGG pathways enrichments analysis was performed using Perseus (Tyanova et al. 2016). The mass spectrometry proteomics data have been deposited to the ProteomeXchange Consortium via the PRIDE partner repository with the dataset identifier PXD024878.

### RNA isolation and quality assessment

*Arabidopsis* total RNA was isolated using a CTAB-based extraction method (Armarego-Marriott et al. 2019) with modifications that are detailed on *protocols.io* (dx.doi.org/10.17504/protocols.io.3f6gjre). The protocol described here was an effective alternative to TRIzol-based extractions for recovering RNA from juvenile de- etiolated tissues. Plant tissues were frozen and ground in liquid nitrogen using mortar and pestle to obtain a fine powder. Ground tissue was kept in safe-lock tubes in liquid nitrogen (or returned to -80C storage) until all samples were processed. Pre-warmed (65°C) extraction buffer (1 mL) [2% Hexadecyltrimethylammonium bromide (CTAB), 2% polyvinylpyrrolidone K 30, 100 mm 2-amino-2-(hydroxymethyl)-1,3-propanediol (TRIS)–HCl, pH 8.0, 25 mm EDTA, 2.0 m NaCl, 0.5 g l−1 spermidine, 2% 2- mercaptoethanol] was added to each tube, mixed well and incubated for 5 mins at 65°C. Then 200 μl of chloroform:IAA (24:1) was added, and spun at 14,000 rcf for 10 mins. The upper aqueous phase was recovered. Later an equal volume of 5 M LiCl was added into the aqueous layer and mixed well. After overnight incubation at -20°C, the RNA pellet was collected by centrifugation at 14 000 rcf for 20 min at 4°C. Then the pellet was washed with 80 % ethanol and dissolved in 200 μl RNase-free water. The quality of RNA samples was assessed with BioAnalyser / LabChip GXII or by visualization of ∼50 ng RNA on a 1% agarose gel. RNA was treated with TURBO DNase (Ambion/ThermoFisher) followed by AMPure cleanup (detailed protocol is available in GitHub repository https://goo.gl/jMXW1F). RNA was quality assessed for RIN scores using a Bioanalyser (Agilent) or a GXII labchip (Perkin Elmer). RNA quantity and purity was assessed using the ND-1000 spectrophotometer (Nano- Drop Technologies). RNA quality was assessed using the LabChip GXII (Perkin-Elmer).

### Library preparation and sequencing

For total RNA-seq, 1 ug of RNA was depleted for rRNA using the Illumina Plant Ribo-Zero kit, following the manufacturer’s protocol (Ribo-Zero Magnetic Kits Guide Rev A and TruSeq Stranded Total RNA Guide Rev E), except all reaction volumes were adjusted by 1/2 (Crisp et al. 2017). Depleted RNA was then purified and fragmented (7 min) using the Illumina Truseq mRNA stranded kit with 1/3 adjusted reaction volumes (Crisp et al. 2017). An optimal cycle number of 8 was used for the amplification of the final libraries. Samples were pooled in equal molar ratios and sequenced on the NextSeq500 (75 bp single-end) at the ACRF Biomolecular Research Facility (Australian National University, ACT, Australia). Raw sequence data is available from GEO (GSE156677).

### Quantification of transcripts

Raw sequencing reads were trimmed to remove adaptor sequences and low-quality base calls (-q 20—length 80—stringency 3) by using *Trim Galore!* (v0.4.1) (Cheng et al. 2016). Then Subsequent read quality was assessed using *FastQC* (v0.10.1) (Andrews, 2010). Transcript quantification was performed using *Kallisto* (Bray et al. 2016) in conjunction with the *Arabidopsis thaliana* Reference Transcript Dataset 2 (AtRTD2_19April2016.gtf), which contains a comprehensive set of transcript sequences (Cao et al. 2017). Code is available on GitHub (https://github.com/dtrain16/NGS-scripts/blob/master/RNA/RNAseq_kallisto_v1.sh).

### Green Index determination

To semi quantitatively assess the greening of seedling during the light to dark transition, photos of the different genotypes were taken and independent cotyledons (n=10) for each genotype and time point were used to determine the Green Index (GI). Moreover, the GI determinations were performed in five independent experiments. GI was performed by recording the RGB values obtained from as the average of a 5x5 pixel area using Affinity Photo software. Then GI was calculated as follows:

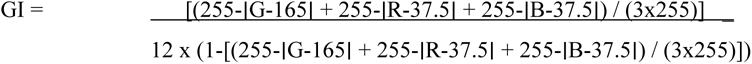

R, G, and B refers to the red, green and blue (R, G, B) of the RGB values and ranges from 0 to 255.

### Correlation analysis between transcriptomic and proteomic data

A correlation between proteomic and transcriptomic data was performed for the subset of plastid proteins IDs that matched the criteria of being differentially enriched in at least three out of four autophagy mutants tested and consistent in at least two out of three time point tested. To do this, the Log2 LFQ intensities were normalized to the average for this subset of proteins and the equivalent normalization of Log2 TPM was done for the transcriptomic data. The normalized Log2 LFQ and Log2 TPM were plotted as scatter plot in R and a Pearson correlation test was performed in Excel to determine the Person correlation coefficient, the *p*- value and its biological significance.

### Chlorophyll fluorescence imaging and photosynthetic parameters

Images of chlorophyll fluorescence were measured using the MAXI-Imaging PAM fluorometer system (Walz, Germany) (Barbagallo et al. 2003). Plants were dark-adapted for 15 min before minimal fluorescence (Fo) was measured. Then, maximal fluorescence (Fm) was measured during 800 ms exposure to a saturating pulse, having a photon flux density (PFD) of 4800 μmol·m^-2^·s^-1^. Thereafter, Fv/Fm, the maximum quantum efficiency of PSII photochemistry, was calculated. Electron transport rate (ETR) parameter was determined immediately after Fo and Fm determination. Twenty measurements were used to describe the light curve for ETR, from 0 to 1076 photosynthetic active radiation (PAR), with a 20 seconds gap between measurements. At least six biological replicates were used per experiment and four independent experiments were combined to obtain the final result.

### Chlorophyll extraction and analysis

Frozen de-etiolating seedlings were ground in a mortar under liquid nitrogen. 20 mg of tissue were transferred into 400 μL of methanol and shaken for two minutes at 30 hz. Samples were centrifuged for two minutes at 14000 g at 4°C. The supernatant was re-extracted with a second aliquot of 400 μL of methanol, shaken for 2 minutes, and debris pelleted before measurement at 652 nm and 665 nm. The formula used to determine the chlorophyll content were: chlorophyll a (μg/mL) = (-8.0962 A652, 1 cm + 16.5169 A665, 1 cm), chlorophyll b (μg/mL) = (27.4405 A652, 1 cm – 12.1688 A665, 1 cm) (Warren 2008).

### Microscopic analysis of chloroplast

Confocal laser scanning microscopy (CLSM) (Nikon A1-Si Confocal Microscope) was used to examine plastids during the photomorphogenesis transition. Chlorophyll auto-fluorescence was used to visualize with filters set at excitation: 488 nm; emission: 640 nm. Images were acquired sequentially or simultaneously where appropriate (Kodama 2016). Resulting regions of interest (ROIs) were selected as 1024x1024 px^2^; area (in px^2^) where a pixel resolution size 0.07 μm/px. A total of 5 biological replicates were used to determine the chloroplast size and 10 biological replicates for counting the chloroplasts.

### Liquid-phase oxygen evolution

Oxygen evolution was determined using a Clark type polarographic oxygen electrode (Oxygraph Plus System). Oxygen electrode were assembled, stabilized for 1h and calibrated every day prior measurements. The liquid phase consisted of 1 mM NaHCO_3_, 10 mM HEPES-KOH, pH 7.5 buffer, which was freshly prepared every day. Seedlings were weighed on an analytical scale and fresh weight (FW) was recorded. A minimum of 6 mg of seedlings was used in every case. Oxygraph temperature was set to 25°C and stirrer speed at 70 rpm. Seedlings were transferred to the Oxygraph chamber containing 2 mL of the above-mentioned buffer, and three phase measurements were taken. The first phase consisted of 3-5 min of measurement under dark condition to determine oxygen consumption (i.e. respiration). After having at least 3 min of linear slope, the second phase was initiated by turning on the light and the oxygen evolution was recorded for at least 5 additional min. A 12V 100W Olympus LG-PS2 Fibre Optic Illuminator was used as source of light to induce photosynthetic activity. After this time, light was turned off again and another 3-5 min were recorded in darkness (third phase). Respiration was calculated as the average of the oxygen consumption determined from the slope of phase 1 and 3 per FW of sample and expressed as nmol O_2_ · min^-1^· g^-1^ FW. Light dependent oxygen production (i.e. photosynthesis) was calculated as the difference between the slope obtained in light condition (phase 2) and darkness (phase 1 and 3), and expressed as nmol O_2_ ·min^-1^·g^-1^ FW. The ratio between these two parameters (photosynthesis/respiration) was also determined to give an indication how photoautotrophic the seedlings were and thus their developmental stage. Values higher than 1 mean the plant produce more oxygen than what it consumes under light condition indicating a more developed state, whereas values lower than 1 indicates the plant consume more oxygen than what it produces indicating a less developed state. The measurements consisted of at least six biological replicates, each biological replicate consisting of multiple seedlings.

## Supplemental Data

**Supplemental Table 1.** Green Index Tukey’s HSD comparison for autophagy and lipid metabolism mutants and WT.

**Figure S1.** Abundance of plastid MRM protein biomarkers in WT during germination (T1) and light to dark transition (T3).

**Figure S2.** Organelles specific protein abundance profiling of WT and autophagy mutants in the dark to light transition

**Figure S3.** Comparison of relative apparent electron transport rate (ETR) plots between the different genotypes and WT.

**Figure S4.** Liquid-phase respiration and photosynthetic activity in different genotypes and WT.

**Figure S5.** Comparison of relative apparent electron transport rate (ETR) plots between the different genotypes and WT

**Figure S6.** Comparison of relative apparent electron transport rate (ETR) plots for WT and autophagy mutants.

## Acknowledgment

We thank the authors referenced in methods for provision of autophagy mutants and especially thank Dr Changcheng Xu from Brookhaven National Laboratory for provision of the series of *sdp1*, *tgd1* and *atg* double mutants reported in Fan et al. (2019) that were used in this study. Dr Brendan O’Leary (The University of Western Australia) is thanked for critical reading of the article. AWY was supported by a UWA Research Training Program International Student Scholarship. This work was support by Australian Research Council funding to AHM and BJP (CE140100008). We acknowledge the Biomolecular Resource Facility at the ANU for performing Illumina sequencing and the provision of computational infrastructure by the National Computational Infrastructure supported under the National Collaborative Research Infrastructure Strategy of the Australian Government.

## Author contributions

AWY and SS performed most of the experiments and data analysis, RF and OD assisted with mass spectrometry and data analysis, DRG and BJP assisted with RNAseq and data analysis, AHM, OD, LL, ES and SS designed the experiments. SS and AHM wrote the paper and all authors contributed to revising it.

## REFERENCES

Armarego-Marriott, Kowalewska, Burgos, Fischer, Thiele, Erban, Strand, Kahlau, Hertle, Kopka, et al (2019) Highly resolved systems biology to dissect the etioplast-to-chloroplast transition in tobacco leaves. Plant Physiol 180:654–681

Avin-Wittenberg (2018) Autophagy and its Role in Plant Abiotic Stress Management. Plant Cell Environ. https://doi.org/10.1111/pce.13404

Avin-Wittenberg, Bajdzienko, Wittenberg, Alseekh, Tohge, Bock, Giavalisco, Fernie (2015) Global analysis of the role of autophagy in cellular metabolism and energy homeostasis in Arabidopsis seedlings under carbon starvation. Plant Cell 27:306–322

Awai, Xu, Tamot, Benning (2006) A phosphatidic acid-binding protein of the chloroplast inner envelope membrane involved in lipid trafficking. Proc Natl Acad Sci U S A 103:10817–10822

Baena-González, Rolland, Thevelein, Sheen (2007) A central integrator of transcription networks in plant stress and energy signalling. Nature 448:938–942

Barbagallo, Oxborough, Pallett, Baker (2003) Rapid, noninvasive screening for perturbations of metabolism and plant growth using chlorophyll fluorescence imaging. Plant Physiol 132:483–493

Bassham (2007) Plant autophagy - more than a starvation response. Curr Opin Plant Biol 10:587–593

Bray, Pimentel, Melsted, Pachter (2016) Near-optimal probabilistic RNA-seq quantification. Nat Biotechnol 34:525–527

Broda, Millar, Van Aken (2018) Mitophagy: A Mechanism for Plant Growth and Survival. Trends Plant Sci 23:434–450

Cai, Li, Liu, Wang, Zhou, Xu, Xiong (2017) COP1 integrates light signals to ROP2 for cell cycle activation. Plant Signal Behav 12:

Calixto, Guo, James, Tzioutziou, Entizne, Panter, Knight, Nimmo, Zhang, Brown (2018) Rapid and dynamic alternative splicing impacts the arabidopsis cold response transcriptome[CC-BY]. Plant Cell 30:1424–1444

Cao, Ni, Zhang, Shi, Xu, Yan, Zhang (2017) Alterations in the proteome of wheat primary roots after wortmannin application during seed germination. Acta Physiol Plant 39:223

Chen, Liao, Qi, Xie, Huang, Tan, Zhai, Yuan, Zhou, Yu, et al (2015) Autophagy contributes to regulation of the hypoxia response during submergence in Arabidopsis thaliana. Autophagy 11:2233–2246

Chen, Liu, Xiong, Sheen, Wu (2018) TOR and RPS6 transmit light signals to enhance protein translation in deetiolating Arabidopsis seedlings. Proc Natl Acad Sci U S A 115:12823– 12828

Cheng, Teo, Krueger, Rock, Koh, Choi, Vogel (2016) Differential dynamics of the mammalian mRNA and protein expression response to misfolding stress. Mol Syst Biol 12:855

Chung, Suttangkakul, Vierstra (2009) The ATG autophagic conjugation system in maize: ATG transcripts and abundance of the ATG8-lipid adduct are regulated by development and nutrient availability. Plant Physiol 149:220–234

Cox, Mann (2008) MaxQuant enables high peptide identification rates, individualized p.p.b.- range mass accuracies and proteome-wide protein quantification. Nat Biotechnol 26:1367–1372

Crisp, Ganguly, Smith, Murray, Estavillo, Searle, Ford, Bogdanović, Lister, Borevitz, et al (2017) Rapid recovery gene downregulation during excess-light stress and recovery in arabidopsis. Plant Cell 29:1836–1863

Cui, Hayashi, Otomo, Mano, Oikawa, Hayashi, Nishimura (2016) Sucrose production mediated by lipid metabolism suppresses the physical interaction of peroxisomes and oil bodies during germination of arabidopsis thaliana. J Biol Chem 291:19734–19745

Di Berardino, Marmagne, Berger, Yoshimoto, Cueff, Chardon, Masclaux-Daubresse, Reisdorf-Cren (2018) Autophagy controls resource allocation and protein storage accumulation in Arabidopsis seeds. J Exp Bot 69:1403–1414

Eastmond (2006) Sugar-dependent1 encodes a patatin domain triacylglycerol lipase that initiates storage oil breakdown in germinating Arabidopsis seeds. Plant Cell 18:665–675

Fan, Yu, Xu (2017) A central role for triacylglycerol in membrane lipid breakdown, fatty acid β-oxidation, and plant survival under extended darkness. Plant Physiol 174:1517– 1530

Fan, Yu, Xu (2019) Dual role for autophagy in lipid metabolismin arabidopsis. Plant Cell 31:1598–1613

Ghillebert, Swinnen, Wen, Vandesteene, Ramon, Norga, Rolland, Winderickx (2011) The AMPK/SNF1/SnRK1 fuel gauge and energy regulator: Structure, function and regulation. FEBS J 278:3978–3990

Guiboileau, Yoshimoto, Soulay, Bataillé, Avice, Masclaux-Daubresse (2012) Autophagy machinery controls nitrogen remobilization at the whole-plant level under both limiting and ample nitrate conditions in Arabidopsis. New Phytol 194:732–740

Guo, Tzioutziou, Stephen, Milne, Calixto, Waugh, Brown, Zhang (2020) 3D RNA-seq: a powerful and flexible tool for rapid and accurate differential expression and alternative splicing analysis of RNA-seq data for biologists. RNA Biol. https://doi.org/10.1080/15476286.2020.1858253

Havé, Luo, Tellier, Balliau, Cueff, Chardon, Zivy, Rajjou, Cacas, Masclaux-Daubresse (2019) Proteomic and lipidomic analyses of the Arabidopsis atg5 autophagy mutant reveal major changes in endoplasmic reticulum and peroxisome metabolisms and in lipid composition. New Phytol 223:1461–1477

Heredia-Martínez, Andrés-Garrido, Martínez-Force, Pérez-Pérez, Crespo (2018) Chloroplast damage induced by the inhibition of fatty acid synthesis triggers autophagy in chlamydomonas. Plant Physiol 178:1112–1129

Hooper, Stevens, Saukkonen, Castleden, Singh, Mann, Fabre, Ito, Deery, Lilley, et al (2017) Multiple marker abundance profiling: combining selected reaction monitoring and data- dependent acquisition for rapid estimation of organelle abundance in subcellular samples. Plant J 92:1202–1217

Ishida, Izumi, Wada, Makino (2014) Roles of autophagy in chloroplast recycling. Biochim Biophys Acta - Bioenerg 1837:512–521

Izumi, Ishida, Nakamura, Hidema (2017) Entire Photodamaged Chloroplasts Are Transported to the Central Vacuole by Autophagy. Plant Cell 29:377–394

Izumi, Wada, Makino, Ishida (2010) The autophagic degradation of chloroplasts via rubisco- containing bodies is specifically linked to leaf carbon status but not nitrogen status in arabidopsis. Plant Physiol 154:1196–1209

Janse van Rensburg, Van den Ende, Signorelli (2019) Autophagy in plants: Both a puppet and a puppet master of sugars. Front Plant Sci 10:

Jiang, Zhu, Wang, Hou (2020) Autophagy-related 2 regulates chlorophyll degradation under abiotic stress conditions in arabidopsis. Int J Mol Sci 21:4515

Kodama (2016) Time gating of chloroplast autofluorescence allows clearer fluorescence imaging in planta. PLoS One 11:e0152484

LaBrant, Barnes, Roston (2018) Lipid transport required to make lipids of photosynthetic membranes. Photosynth Res 138:345–360

Lai, Wang, Zheng, Fan, Chen (2011) A critical role of autophagy in plant resistance to necrotrophic fungal pathogens. Plant J 66:953–968

Lastdrager, Hanson, Smeekens (2014) Sugar signals and the control of plant growth and development. J Exp Bot 65:799–807

Law, Narsai, Whelan (2014) Mitochondrial biogenesis in plants during seed germination. Mitochondrion 19:214–221

Li-Beisson, Shorrosh, Beisson, Andersson, Arondel, Bates, Baud, Bird, DeBono, Durrett, et al (2013) Acyl-Lipid Metabolism. In: The Arabidopsis Book. p e0161

Li, Chung, Vierstra (2014) AUTOPHAGY-RELATED11 plays a critical role in general autophagy- and senescence-induced mitophagy in Arabidopsis. Plant Cell 26:788–807

Li, Gao, Benning, Sharkey (2012) Characterization of photosynthesis in Arabidopsis ER-to- plastid lipid trafficking mutants. Photosynth Res 112:49–61

Li, Yu (2018) Chloroplast galactolipids: The link between photosynthesis, chloroplast shape, jasmonates, phosphate starvation and freezing tolerance. Plant Cell Physiol 59:1128– 1134

Luo, Zhou, Masclaux-Daubresse, Wang, Wang, Zheng (2019) Morphological and physiological responses to contrasting nitrogen regimes in Populus cathayana is linked to resources allocation and carbon/nitrogen partition. Environ Exp Bot 162:247–255

Ma, Liang, Zhao, Wang, Ma, Mai, Andrade, Zeng, Grujic, Jiang, et al (2021) Article Friendly mediates membrane depolarization- induced mitophagy in Arabidopsis. *Curr Biol* doi.org/10.1016/j.cub.2021.02.034.

Marshall, Vierstra (2018) Autophagy: The Master of Bulk and Selective Recycling. Annu Rev Plant Biol 69:173–208

Masclaux-Daubresse, Chen, Havé (2017) Regulation of nutrient recycling via autophagy. Curr Opin Plant Biol 39:8–17

McLoughlin, Augustine, Marshall, Li, Kirkpatrick, Otegui, Vierstra (2018) Maize multi-omics reveal roles for autophagic recycling in proteome remodelling and lipid turnover. Nat Plants 4:1056–1070

McLoughlin, Marshall, Ding, Chatt, Kirkpatrick, Augustine, Li, Otegui, Vierstra (2020) Autophagy plays prominent roles in amino acid, nucleotide, and carbohydrate metabolism during fixed-carbon starvation in maize. Plant Cell 32:2699–2724

Michaud, Jouhet (2019) Lipid trafficking at membrane contact sites during plant development and stress response. Front Plant Sci 10:1–10

Minina, Moschou, Vetukuri, Sanchez-Vera, Cardoso, Liu, Elander, Dalman, Beganovic, Lindberg Yilmaz, et al (2018) Transcriptional stimulation of rate-limiting components of the autophagic pathway improves plant fitness. J Exp Bot 69:1415–1432

Nukarinen, Ngele, Pedrotti, Wurzinger, Mair, Landgraf, Börnke, Hanson, Teige, Baena-Gonzalez, et al (2016) Quantitative phosphoproteomics reveals the role of the AMPK plant ortholog SnRK1 as a metabolic master regulator under energy deprivation. Sci Rep 6:1–19

O’Leary, Oh, Lee, Millar (2020). Metabolite regulatory interactions control plant respiratory metabolism via Target of Rapamycin (TOR) Kinase activation. Plant Cell. doi.org/10.1105/tpc.19.00157

Pfeiffer, Janocha, Dong, Medzihradszky, Schöne, Daum, Suzaki, Forner, Langenecker, Rempel, et al (2016) Integration of light and metabolic signals for stem cell activation at the shoot apical meristem. Elife 5:

Pogson, Ganguly, Albrecht-Borth (2015) Insights into chloroplast biogenesis and development. Biochim Biophys Acta - Bioenerg 1847:1017–1024

Ritchie, Phipson, Wu, Hu, Law, Shi, Smyth (2015) Limma powers differential expression analyses for RNA-sequencing and microarray studies. Nucleic Acids Res 43:e47

Robinson, McCarthy, Smyth (2010) edgeR: A Bioconductor package for differential expression analysis of digital gene expression data. Bioinformatics 26:139–140

Robinson, Oshlack (2010) A scaling normalization method for differential expression analysis of RNA-seq data. Genome Biol 11:R25

Rudowska, Gieczewska, Mazur, Garstka, Mostowska (2012) Chloroplast biogenesis - Correlation between structure and function. Biochim Biophys Acta - Bioenerg 1817:1380–1387

Schnurr, Shockey, De Boer, Browse (2002) Fatty acid export from the chloroplast. Molecular characterization of a major plastidial acyl-coenzyme A synthetase from Arabidopsis. Plant Physiol 129:1700–1709

Signorelli, Tarkowski, Van den Ende, Bassham (2019) Linking Autophagy to Abiotic and Biotic Stress Responses. Trends Plant Sci 24:

Soneson, Love, Robinson (2016) Differential analyses for RNA-seq: Transcript-level estimates improve gene-level inferences [version 2; referees: 2 approved]. F1000Research 4:1521

Thazar-Poulot, Miquel, Fobis-Loisy, Gaude (2015) Peroxisome extensions deliver the Arabidopsis SDP1 lipase to oil bodies. Proc Natl Acad Sci U S A 112:4158–4163

Tyanova, Temu, Cox (2016) The MaxQuant computational platform for mass spectrometry-based shotgun proteomics. Nat Protoc 11:2301–2319

Warren (2008) Rapid measurement of chlorophylls with a microplate reader. J Plant Nutr 31:1321–1332

Waters, Langdale (2009) The making of a chloroplast. EMBO J 28:2861–2873

Wessel, Flügge (1984) A method for the quantitative recovery of protein in dilute solution in the presence of detergents and lipids. Anal Biochem 138:141–143

Xu, Fan, Froehlich, Awai, Benning (2005) Mutation of the TGD1 chloroplast envelope protein affects phosphatidate metabolism in Arabidopsis. Plant Cell 17:3094–3110

Xu, Fan, Riekhof, Froehlich, Benning (2003) A permease-like protein involved in ER to thylakoid lipid transfer in Arabidopsis. EMBO J 22:2370–2379

Xu, Moellering, Muthan, Fan, Benning (2010) Lipid transport mediated by arabidopsis TGD proteins is unidirectional from the endoplasmic reticulum to the plastid. Plant Cell Physiol 51:1019–1028

Yoshimoto, Jikumaru, Kamiya, Kusano, Consonni, Panstruga, Ohsumi, Shirasu (2009) Autophagy negatively regulates cell death by controlling NPR1-dependent salicylic acid signaling during senescence and the innate immune response in Arabidopsis. Plant Cell 21:2914–27

Young, Bartel (2016) Pexophagy and peroxisomal protein turnover in plants. Biochim Biophys Acta - Mol Cell Res 1863:999–1005

Zhuang, Chung, Cui, Lin, Gao, Kang, Jiang (2017) ATG9 regulates autophagosome progression from the endoplasmic reticulum in Arabidopsis. Proc Natl Acad Sci 114:E426–E435

